# Intestinal microbiota programming of alveolar macrophages influences severity of respiratory viral infection

**DOI:** 10.1101/2023.09.21.558814

**Authors:** Vu L. Ngo, Carolin M. Lieber, Hae-ji Kang, Kaori Sakamoto, Michal Kuczma, Richard K. Plemper, Andrew T. Gewirtz

**Affiliations:** Center for Translational Antiviral Research, Georgia State University Institute for Biomedical Sciences, Atlanta GA 30303, USA; Department of Pathology, University of Georgia College of Veterinary Science, Athens GA 30602, USA

**Author notes:** These authors contributed equally to this work (Vu L. Ngo, Carolin M. Lieber). Denotes co-senior and co-corresponding authors (Richard K. Plemper, Andrew T. Gewirtz).

## Abstract

Susceptibility to respiratory virus infections (RVIs) varies widely across individuals. Because the gut microbiome impacts immune function, we investigated the influence of intestinal microbiota composition on RVI and determined that segmented filamentous bacteria (SFB), naturally acquired or exogenously administered, protected mice against influenza virus (IAV) infection. Such protection, which also applied to respiratory syncytial virus and SARS-CoV-2, was independent of interferon and adaptive immunity but required basally resident alveolar macrophages (AM). In SFB-negative mice, AM were quickly depleted as RVI progressed. In contrast, AM from SFB-colonized mice were intrinsically altered to resist IAV-induced depletion and inflammatory signaling. Yet, AM from SFB-colonized mice were not quiescent. Rather, they directly disabled IAV via enhanced complement production and phagocytosis. Accordingly, transfer of SFB-transformed AM into SFB-free hosts recapitulated SFB-mediated protection against IAV. These findings uncover complex interactions that mechanistically link the intestinal microbiota with AM functionality and RVI severity.

**One sentence summary:** Intestinal segmented filamentous bacteria reprogram alveolar macrophages promoting nonphlogistic defense against respiratory viruses.

## INTRODUCTION

Outcomes following exposure to respiratory viruses, such as influenza A viruses (IAV), respiratory syncytial virus (RSV), and severe acute respiratory syndrome coronavirus 2 (SARS-CoV-2), vary widely amongst individuals, ranging from asymptomatic infection to severe lung pathology and/or death. While determinants of such heterogeneity to RVI are likely numerous and complex, we hypothesized that one important factor is gut microbiota composition ^1^, which is now appreciated to have broad influence over a range of chronic inflammatory diseases and the immune responses with which such diseases are associated ^2^. For example, studies comparing mice captured in the wild to those bred for generations in well-controlled vivaria revealed vast differences in their microbiomes that resulted in wild mice having highly and broadly activated immune systems and being relatively resistant to IAV ^3,4^. Yet, differences in specific microbial species that can impact RVI outcomes, and how they might do so, have not been well defined. Hence, we studied mice raised in vivaria with discrete defined microbiome differences and, subsequently, mice differing in only the presence or absence of a single common species, namely segmented filamentous bacteria (SFB). We found that gut SFB dictated the phenotype of alveolar macrophages and, consequently, modulated outcomes of RVI.

## RESULTS

### SFB protects mice from IAV infection

Appreciation of the potential of microbiota to broadly impact health led one commercial rodent supplier to offer “excluded flora (EF)” mice, which were bred in a restricted vivarium that assured absence of a discrete panel of disease-modulating commensal microbes, which might or might not otherwise be present in colonies of “specific pathogen-free (SPF)” mice commonly used in biomedical research ^5^. We used such mice as a starting point to probe the influence of microbiota composition on RVI. Specifically, we compared proneness of SPF and EF mice to IAV infection following intranasal administration of 2009 pandemic A/CA/07/2009 (H1N1), herein referred to as CA09, which readily infects mice without need for species adaptation and recapitulates major clinical features of human disease in the animal model ^6^. Viral titers in the lung were measured 4 days post-infection (dpi), a time point at which disease symptoms, including mortality, could be assessed although experiments explicitly designed to measure survival were allowed to proceed until 14 dpi, when infectious particles were no longer detectable. Compared to EF mice, SPF mice exhibited significantly reduced lung virus loads. Furthermore, SPF mice did not show hypothermia and weight loss observed in EF mice (**Figure 1A**). Additionally, some mice of the EF mice required euthanasia. These results suggested that one or more of the microbes specifically excluded from EF mice might confer protection against IAV.

**Figure 1.**
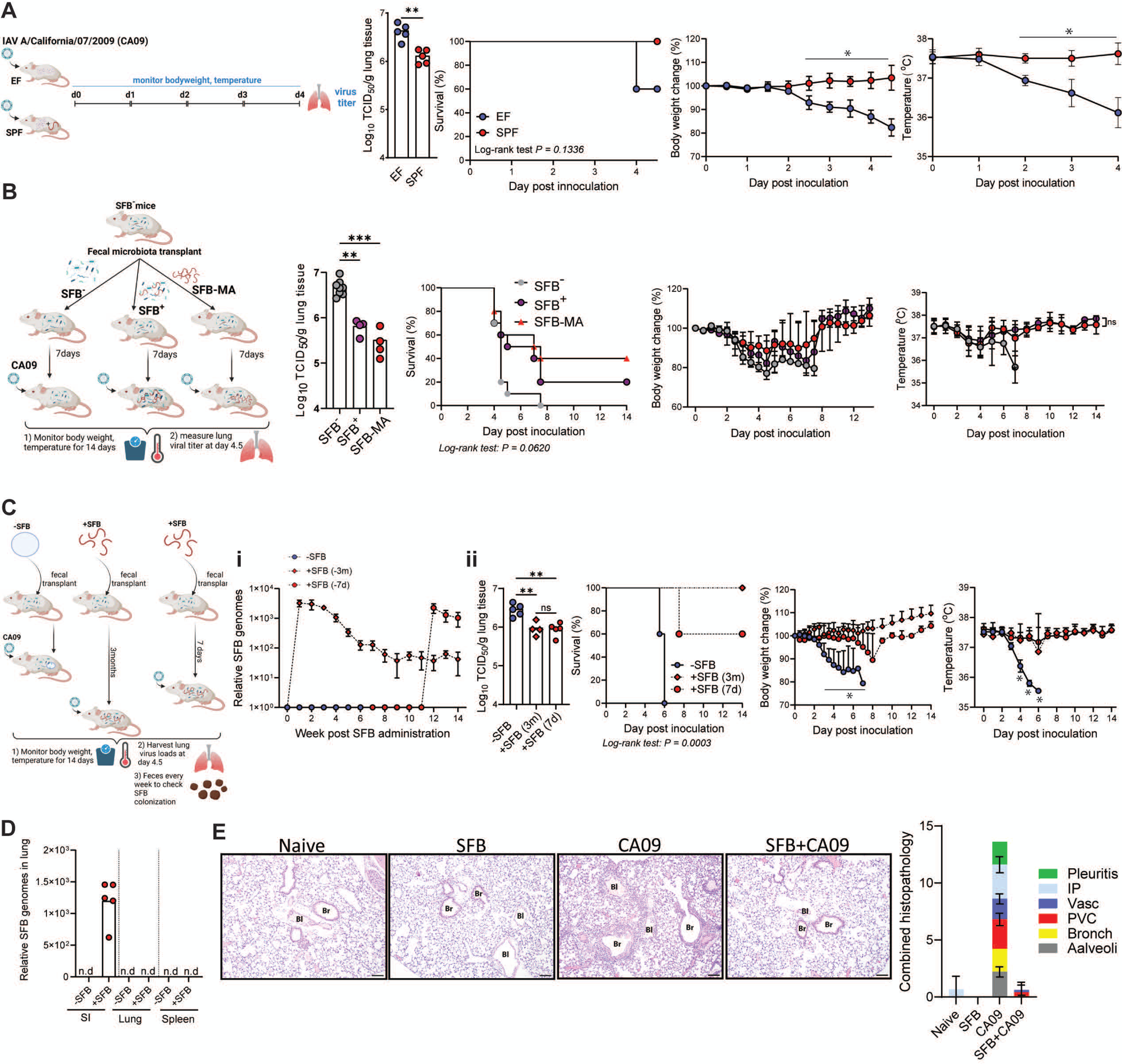
Colonization of the intestine with SFB protected mice from influenza virus infection. **(A)** EF and SPF mice were inoculated with A/CA/07/2009 (H1N1) influenza virus (CA09) as schematized. Lung viral titers, survival, bodyweight, and temperature were monitored. **(B)** As schematized, conventionally colonized SFB^-^ mice received feces that lacked SFB (SFB^-^), contained SFB (SFB^+^), or contained SFB but no other bacteria (SFB-MA). Seven days post-transplant, mice were inoculated with CA09. Mice were euthanized 4.5-days later or monitored daily for survival, body weight, and temperature for 2 weeks. Statistical analyses were performed with SFB^-^ as a control group. **(C)** SFB^-^ mice were administered SFB 3 months or 7 days prior to CA09 inoculation. **(i)** qRT-PCR analysis of SFB colonization level over the course of 14 weeks. **(ii)** Mice were euthanized 4.5-days later or monitored daily for survival, body weight, and temperature for 2 weeks. **(D)** qRT-PCR analysis of SFB level in the small intestine, lungs, and spleens after 7 days colonization. **(E)** SFB^-^ mice were untreated (naive) or administrated SFB. Seven days later, mice were inoculated (or not) with CA09. Mice were euthanized 4.5 days later. H&E staining lung sections scored for pleuritis, bronchiolitis, alveolitis, vasculitis, interstitial pneumonia (IP), and perivascular cuffing (PVC). All experiments n = 5 mice per group. Data is representative of two to three independent experiments. yielding an identical pattern of results. Statistical analyses of survival, bodyweight change, and temperature of (B) and (C) were conducted by comparing “SFB-“or “-SFB” group to all other groups. Statistical analysis; Viral titer: One-way ANOVA or student *t* test. survival: Log-rank Mantel-Cox test. Body weights and temperature: Two-way ANOVA. *p<0.05, **p<0.01, ***p<0.001, ns not significant.

Of the panel of organisms known to be absent in EF mice, segmented filamentous bacteria (SFB) stood out as a potential modulator of IAV proneness in that SFB, despite being a strict anaerobe and, thus, restricted to the intestinal luminal surface, is known to systemically impact T-lymphocytes ^7,8^. Furthermore, SFB is a major contributor to, albeit not the sole mediator of, spontaneous resistance of mice to rotavirus (RV), an intestinal pathogen, that arose in some mouse colonies ^9^. SFB is very challenging to culture but can be isolated, maintained, and studied by administering it via fecal microbial transplantation (FMT). Hence, mice lacking SFB but otherwise carrying a complex SPF microbiota, herein referred to as SFB^-^ mice, were administered a FMT from other SFB^-^ mice, mice containing SFB as one natural component of their SPF microbiota, or germ-free mice that had been monoassociated (MA) with SFB. Such mice were inoculated with CA09 one-week post-FMT and euthanized 4d later to measure lung viral titers or monitored for 14 days to assess survival. Mice receiving FMT from donors naturally colonized with SFB exhibited IAV titer reductions of approximately one order of magnitude compared to animals that lacked SFB and, concomitantly, clinical signs of IAV infection were ameliorated (**Figure 1B**). FMT from SFB-MA mice fully recapitulated such protection while administering feces from germ-free mice was without effect (**Figure S1A).** Thus, administering SFB^-^ mice a FMT from SFB-MA mice, hereafter referred to as administering SFB, recapitulated the differential proneness of EF and SPF mice to IAV infection, thus providing a tractable model to investigate how the presence of this specific gut microbiota constituent, within a complex microbial ecosystem, reduced IAV infection in the lung.

SFB-mediated protection against IAV infection was evident, albeit seemingly not maximal, two days post SFB administration and was undiminished three months later (longest time tested) despite gut SFB levels having declined by over 60-fold from peak to a stable plateau by this time (**Figure 1C and Figure S1B**). In accord with its relative abundance not being a critical variable, SFB colonized 24-week-old mice at 10-fold lower levels than 5-week-old mice but still protected them against IAV infection **(Figure S1C)**. Consistent with previous observations that SFB only resides in the gut lumen or attached to the apical epithelial gut surface ^10^, SFB was readily detected by PCR in the ileum, but was not found in the lung or spleen (**Figure 1D**). Histopathologic examination of lung tissue did not detect any impact of SFB itself on the lung but revealed a striking suppression of IAV-induced lung pathology (**Figure 1E**). Such protection of lung tissue was potentially attributable to the 1-log reduction in viral titers conferred by SFB colonization but also hinted at the possibility that SFB had reduced IAV-induced inflammatory signaling.

### SFB-mediated protection against IAV is not explained by known candidate mechanisms

SFB-mediated lasting protection against IAV contrasted with its transient protection against RV ^9^, suggesting that the former might be mediated by adaptive immunity. However, SFB-mediated protection against IAV was fully maintained in *Rag1^-/-^*mice, which lack most T and B lymphocytes, arguing against this notion (**Figure 2A**). Innate antiviral immunity typically involves interferon and can involve innate lymphoid cells (ILC) and/or IL-22 and IL-17. Yet, SFB-mediated protection against IAV was also maintained in *Ifnar^-/-^* mice, which lack type I interferon signaling, *Rag1^-/-^-Il2rg^-/-^*, which lack type 3 ILC, and upon antibody-mediated neutralization of IL-17 and IL-22 (**Figure 2B-D**). SFB-mediated protection against IAV was furthermore maintained in *Stat1^-/-^* animals, which are broadly compromised in IFN signaling (**Figure 2E**). These results argue against a role for any of these candidate host mechanisms in mediating SFB’s protection against IAV infection. We also considered a candidate microbial mechanism. Specifically, we hypothesized that the protection against IAV conferred by SFB might reflect that it induced an array of complex changes in other taxa that might then broadly impact innate immunity. However, SFB’s protection was maintained in Altered Schaedler flora (ASF) mice, which harbor only a minimal 8-species microbial community ^11^, thus arguing against this possibility.

**Figure 2.**
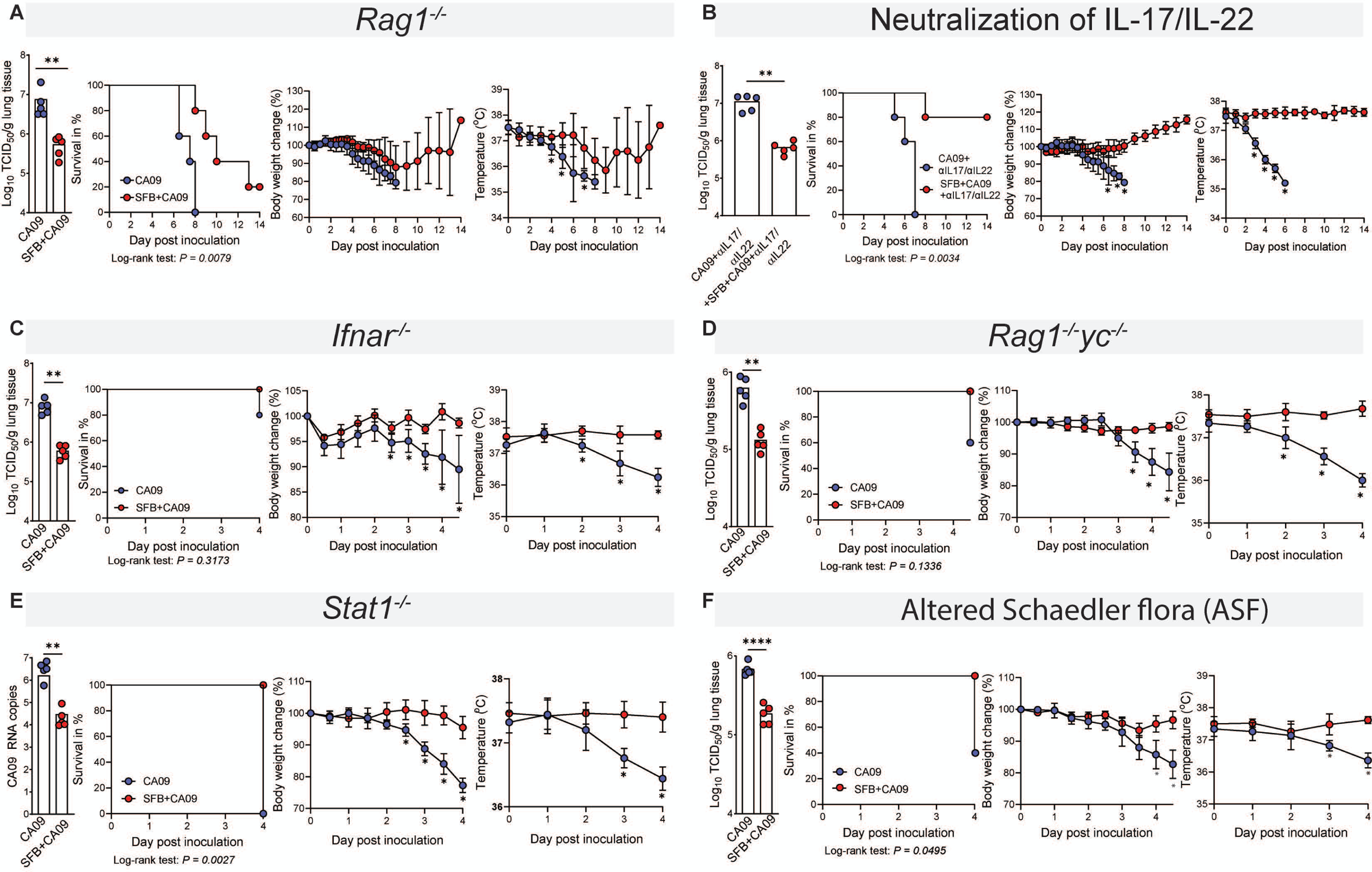
SFB mediated protection against IAV was maintained in an array of knockout mice and did not require complex gut microbiota. Lung viral titer, survival rate, body weight change, and temperature of: SFB-negative mice were colonized with SFB for 7 days then inoculated with CA09. **(A)** *rag1^-/-^* mice **(B)** C57BL/6 mice received neutralization of IL-17/IL-22 monoclonal antibodies every other day throughout the experiment **(C)** *Ifnar^-/-^* mice **(D)** *rag1^-/-^yc^-/-^* mice **(E)** *stat1^-/-^* mice **(F)** Altered Schaedler flora (ASF) mice. All experiments n = 5 mice per group. Statistical analysis; Viral titer: student *t* test, Survival: Log-rank Mantel-Cox test. Body weights and temperature: Two-way ANOVA. **p<0.01, ***p<0.001.

### SFB-mediated protection against IAV, RSV and SARS-CoV-2 is associated with preservation of resident lung phagocytes

That candidate mechanisms were not experimentally substantiated led us to utilize less targeted approaches to investigate how SFB gut colonization protected mice against IAV. First, we performed transcriptomic profiling of whole lungs harvested from mice that were untreated (naïve) or administered SFB, CA09, or a combination thereof. The results of this RNA-seq analysis were presented as a heat map, showing 4-5 individual mice per condition, of all genes whose expression was significantly altered in response to any of the three treatments (**Figure 3A**). It illustrated that SFB, by itself, had almost no impact on lung gene expression, whereas, as expected, IAV infection, by itself, markedly remodeled the lung transcriptome. Strikingly, such IAV-induced changes in gene expression were absent in SFB-colonized mice, indicating that the presence of SFB in the gut broadly ameliorated IAV’s impact on lung tissue. The majority of genes induced by CA09 were related to immunity and inflammation and thus accorded with the view that much of the lung pathology in IAV infection is driven by the immune system ^12^ and, furthermore, the notion that SFB had dampened such responses.

**Figure 3.**
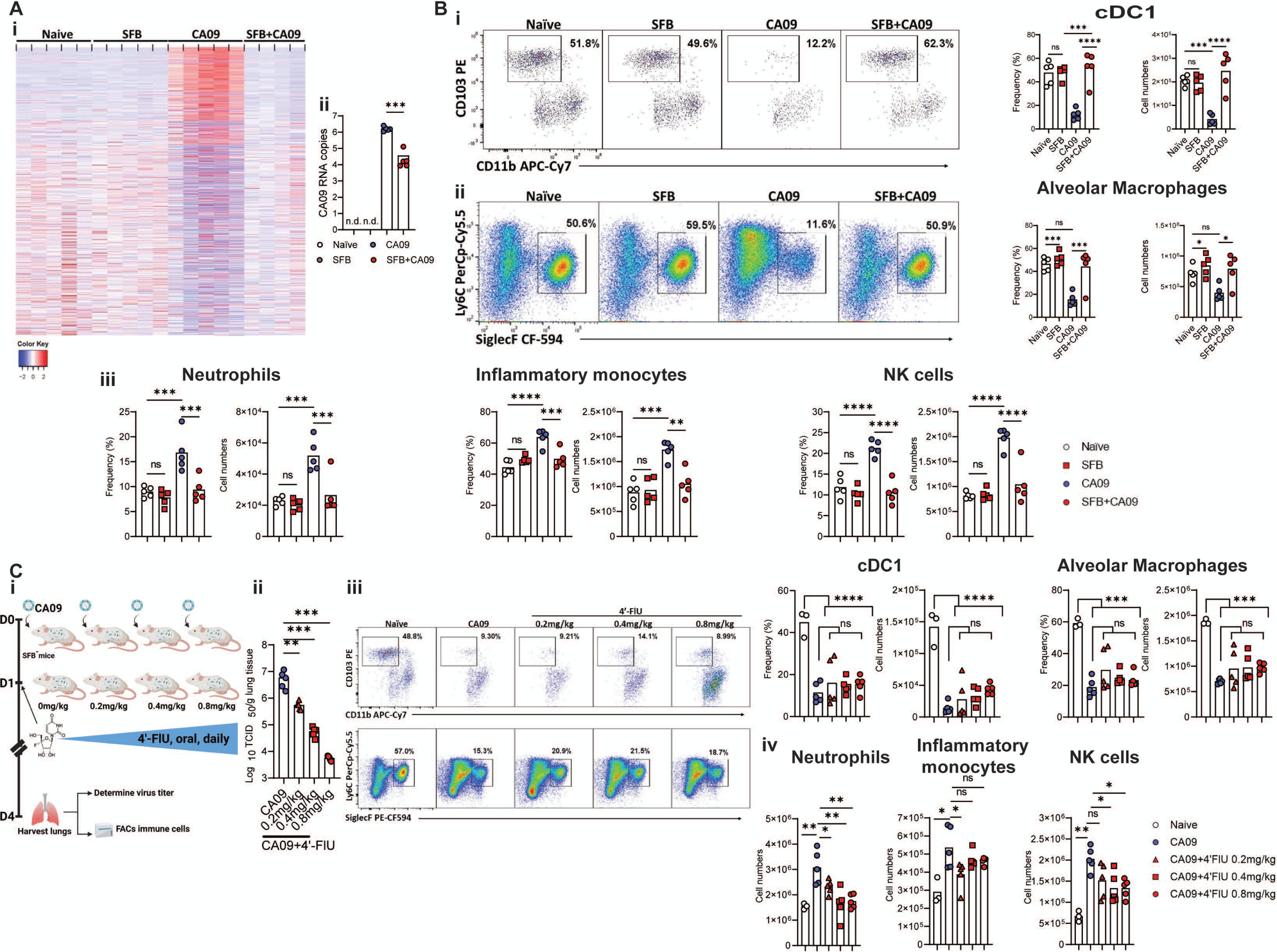
SFB’s protection against IAV infection associated with preservation of resident lung phagocytes. **(A)** SFB^-^ mice were untreated (naïve) or SFB was administered. Seven days later, mice were inoculated (or not) with CA09. Mice were euthanized 4.5-days later. **(i)** Whole lung gene expression assayed by RNA-seq and data are displayed as heatmap of genes that are significantly altered by 2-fold in all treatment groups relative to the naïve group. **(ii)** Quantitation of CA09 level in these extracts by qRT-PCR. **(B)** SFB^-^ mice were untreated (naïve) or SFB was administered. Seven days later, mice were inoculated (or not) with CA09. Mice were euthanized 4.5-days later, and lung assayed by flow cytometry. **(i & ii)** Analysis of dendritic cells and alveolar macrophage; representative flow plot and frequencies and cell numbers **(i)** cDC1. **(ii)** alveolar macrophages. **(iii)** Frequencies and cell numbers of lung neutrophils, inflammatory monocytes, and NK cells. **(C) (i)** Experimental schema: SFB^-^ mice colonized with SFB, inoculated with CA09, and treated with 4’-FlU at indicated doses. On day 4.5 post virus inoculation, mice were euthanized, and lungs were harvested to determine virus titers and FACs analysis. **(ii)** lung viral titer. **(iii)** FACs plot graphical representation, frequencies, and cell numbers of lung cDC1, and AMs. **(iv)** Cell numbers of lung neutrophils, inflammatory monocytes, and NK cells. All experiment n = 4-5 per group. Data is representative of two to three independent experiments (B and C), yielding an identical pattern of results. Statistical analysis: One-way ANOVA. *p<0.05, **p<0.01, ***p<0.001, ****p<0.0001, ns not significant.

We next used flow cytometry to examine the impact of SFB and IAV on lung populations of immune cells, focusing on innate cells given that our results from *Rag1*^-/-^ mice had suggested lymphocytes were not required for SFB-mediated protection against IAV. Consistent with other studies, intranasal administration of CA09, by itself, induced a stark depletion of basally-resident CD103^+^ dendritic cells (cDC1) and alveolar macrophages (AM), which was followed by increased abundance of a panel of inflammatory cells including neutrophils, monocytes, and natural killer cells ^13^. SFB, by itself, had only modest impacts on levels of lung leukocytes, slightly increasing abundance of AM, but wholly prevented the changes induced by CA09 (**Figure 3B and Figure S2)**. Preventing these CA09-induced changes in lung cellularity and disease was not SFB strain-specific as similar protective effects resulted from colonization of the gut by SFB strains isolated in 3 different continents **(Figure S3)**. Nor were SFB impacts specific to IAV. Rather, SFB colonization also reduced viral loads and prevented changes in lung cellularity, and reduced pro-inflammatory gene expression, following inoculation of mice with respiratory syncytial virus (RSV) and mouse-adapted SARS-CoV-2 **(Figure S4)**, which, irrespective of SFB, did not cause clinical signs of disease.

To investigate the extent to which SFB-mediated prevention of IAV-induced changes in lung cellularity simply reflected lower IAV loads, we pharmacologically restricted IAV replication via 4’-fluorouridine (4’-FIU), a broad-spectrum nucleoside analog inhibitor that we recently reported reduces viral titers and disease symptoms up to 60 hours following administration of IAV, RSV, and SARS-CoV-2 ^14,15^. SFB^-^ mice received 4’-FIU via oral gavage at different dose levels in a once daily (q.d.) regimen beginning 24 hours post-inoculation. At this time, IAV levels and immune cell populations were similar in SFB^-^ and SFB**-**administered mice **(Figure S2C)**, which we hereafter refer to as SFB**^+^** mice. As expected, administration of 4’-FIU to SFB^-^ mice dose-dependently suppressed IAV replication to a similar or greater extent than SFB (**Figure 3C**). Like SFB, 4’-FIU administration prevented the CA09-induced appearance of inflammatory cells in the lung. However, even the highest dose of 4’-FIU, which lowered IAV titers by over three orders of magnitude and markedly reduced inflammatory cell recruitment, did not mitigate CA09-induced ablation of cDC1 or AM. This result suggested that the persistence of AM and cDC1 in SFB**^+^** mice was not entirely a consequence of reduced viral loads and/or inflammation but rather implied that maintenance of these cells may contribute to suppression of IAV.

### Alveolar Macrophages are required for SFB-mediated protection against IAV

Potential roles for cDC1 and AM in SFB-mediated protection against IAV were tested in mice harboring deficiencies in these cells. We observed that SFB-mediated protection against IAV was maintained in *Batf3^-/-^* mice, which wholly lack cDC1 ^16^, but was greatly reduced in *Csf2^-/-^*mice, which have a 70% reduction in basal AM levels ^17^ **(Figure 4 A&B)**. The role of AM was further examined by depleting these cells with clodronate liposomes ^18^, which we verified efficiently removed AM but did not alter levels of other lung innate immune cells (**Figure 4Ci**). Depletion of AM, by itself, did not significantly impact IAV titers or clinical signs of disease but eliminated the ability of SFB to lower lung IAV load, ameliorate hypothermia and weight loss, and prevent death (**Figure 4Cii**). These results suggested that SFB-mediated impacts on AM were germane to its protection against RVI. SFB-mediated maintenance of AM levels could conceivably have reflected increased generation of these cells, which can occur from monocyte progenitors migrating to the lung or local self-renewal ^19^, or that SFB colonization had resulted in AM resisting death upon RVI. SFB’s prevention of AM depletion following CA09 inoculation associated with prevention of IAV-induced caspase-3 activation (**Figure 4D**), which marks AM commitment to cell death ^20^, and increased expression of CD71 (**Figure 4E**), which associates with enhanced phagocytosis and survival ^21,22^. Furthermore, SFB colonization increased AM expression of proliferation marker Ki-67 (**Figure 4F**). Collectively, these results suggested that SFB colonization may have increased AM self-renewal and, furthermore, “emboldened” AM to resist IAV-induced depletion, resulting in maintenance of a robust AM population following IAV infection.

**Figure 4.**
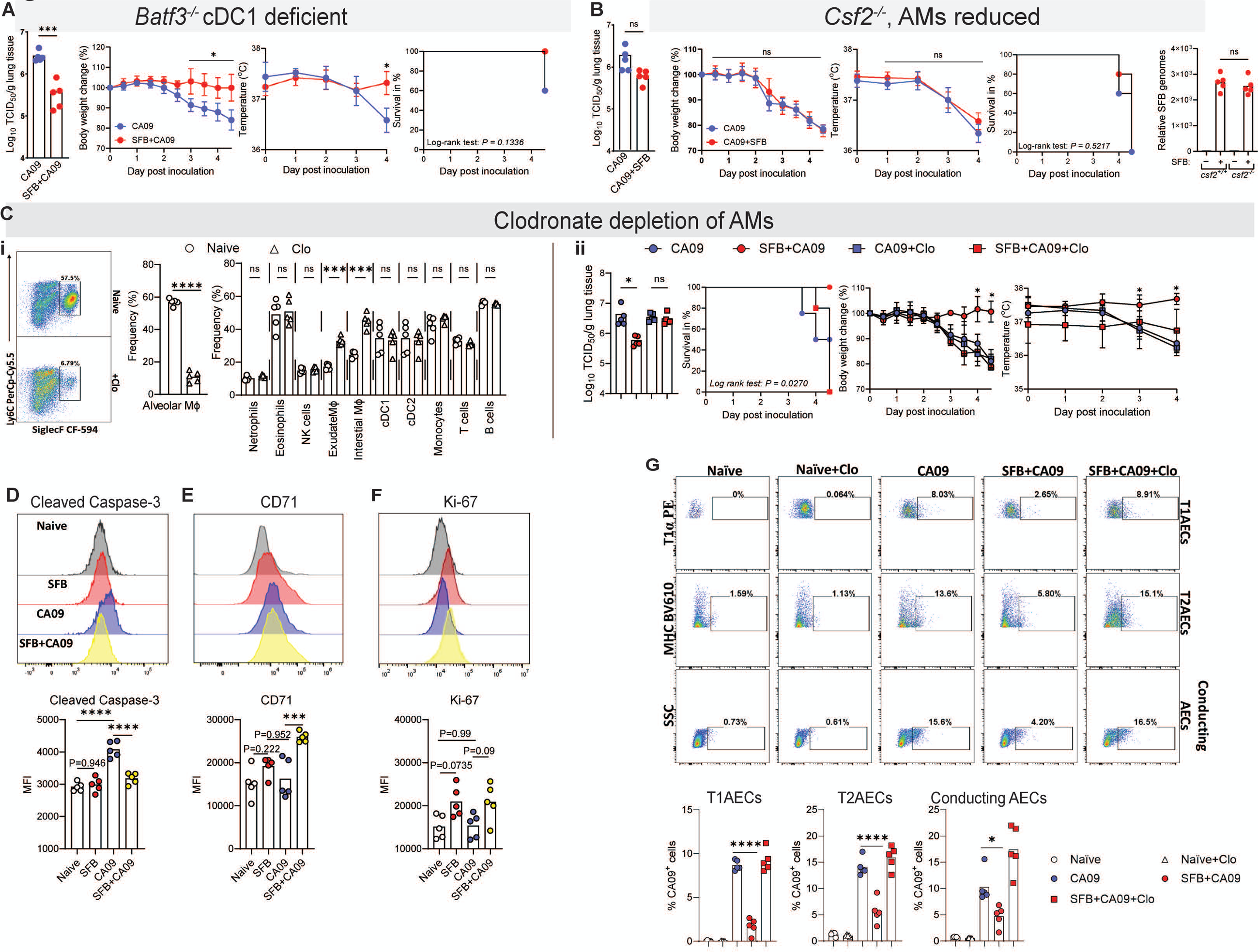
SFB-mediated protection against IAV infection and resulting disease required alveolar macrophages. **(A-B) (A)** *batf3^-/-^* and **(B)** *csf2^-/-^* mice were administered SFB and then, 7 days later, inoculated with CA09. Lung viral titer, body weight change, temperature, and survival were monitored. **(C) (i-ii)** SFB^-^ mice were orally administered with clodronate liposomes one day before mice were euthanized. FACS analysis to determine the AMs depletion efficacy after clodronate treatment. **(i)** FACS graphical representation of lung AMs, frequency of lung AMs and others immune cells after clodronate treatment. **(ii)** SFB^-^ mice were colonized with SFB for 7 days, and clodronate liposomes were orally administered to mice one day before CA09 inoculation. Mice were euthanized on day 4 post-virus inoculation. Lung viral titer, survival rate, body weight, temperature was measured. Statistical analyses of survival, bodyweight change, and temperature were conducted by comparing “SFB+CA09” group to all other groups. **(D)** SFB^-^ mice were untreated (naïve) or SFB was administered. Seven days later, mice were inoculated (or not) with CA09. Mice were euthanized 4.5-days later. Level of cleaved caspase-3 of alveolar macrophages in lung were measured with FACS. **(E-F)** SFB^-^ mice were untreated or treated as in (D). The level of **(E)** CD71 and **(F)** Ki-67 expression of alveolar macrophages in the lungs were measured with FACS. **(G)** SFB^-^ mice treated as described in (C) FACS plot graphical representation, and frequencies CA09 level in T1ACEs, T2AECs, and Conducting AECs were assayed by flow cytometry. All experiment n = 5 mice per group. Data is representative of two to three independent experiments (B-F), yielding an identical pattern of results. Statistical analysis; Viral titer, relative SFB genomes, MFI, frequencies/cell numbers of immune cell data: One-way ANOVA or student *t* test. Survival data Log-rank Mantel-Cox. Body weights, temperature: Two-way ANOVA. *p<0.05, ***p<0.001, ****p<0.0001, ns not significant.

### SFB’s preservation of AM enables their sustained protection of EC from IAV infection

Airway epithelial cells (AEC) are primary replication sites of respiratory viruses, including IAV, RSV, and SARS-CoV-2. Specifically, in the case of highly pathogenic H1N1 IAV, ciliated AEC are the predominant host for virus replication and generation of progeny virions ^23^. AM are known to protect lung AEC from IAV infection, particularly at early time points following IAV-infection, i.e. prior to their depletion ^24^. Thus, we predicted that SFB’s preservation of AM following CA09 inoculation might result in lower viral loads in AEC in SFB^+^ mice. Indeed, the percentage of lung AEC, type I and II, as well as conducting AECs, displaying detectable levels of a recombinant CA09 that encodes GFP, by flow cytometry, was significantly reduced in SFB^+^ mice 4 days post-inoculation (**Figure 4G and Figure S5)**. Depletion of AM by clodronate liposomes in SFB^+^ mice increased IAV levels to that of SFB^-^ mice in all three lung AEC types. These results indicate that SFB-induced AM persistence following IAV infection enables sustained AM-mediated protection of EC, likely contributing to lower IAV burden and ameliorated pathology.

### Persistence of SFB^+^ AM upon RVI is cell-intrinsic

Persistence of AM in CA09-infected SFB^+^ mice was potentially attributable to SFB altering the lung environment and/or reflect that AM themselves had been intrinsically transformed to better withstand IAV challenge. To differentiate between these possibilities, we adoptively transferred FACS-sorted AM from SFB**^+^** and SFB^-^ mice to SFB^-^ mice using donor mice with a CD45.1 subtype, thereby enabling distinction of endogenous and transplanted cells (**Figure 5A**). The degree to which AM harvested from SFB**^+^** and SFB^-^ mice, hereafter referred to as SFB**^+^** and SFB^-^ AM, engrafted following transplant was similar, comprising about 15% of the total AM in their new host. CA09 infection did not significantly alter proportions of endogenous vs engrafted AM in recipients of SFB^-^ AM but led to SFB^+^ AM (i.e. engrafted) becoming their host’s majority AM population. This shift in AM proportions reflected that endogenous AM in both recipient groups as well as engrafted SFB^-^ AM were all depleted by CA09 infection, whereas engrafted SFB**^+^** AM fully resisted CA09-induced depletion and, concomitantly, lacked Caspse-3 activation, despite having been transplanted into an SFB-free host. Thus, colonization of the gut with SFB intrinsically altered AM to withstand IAV-induced depletion.

**Figure 5.**
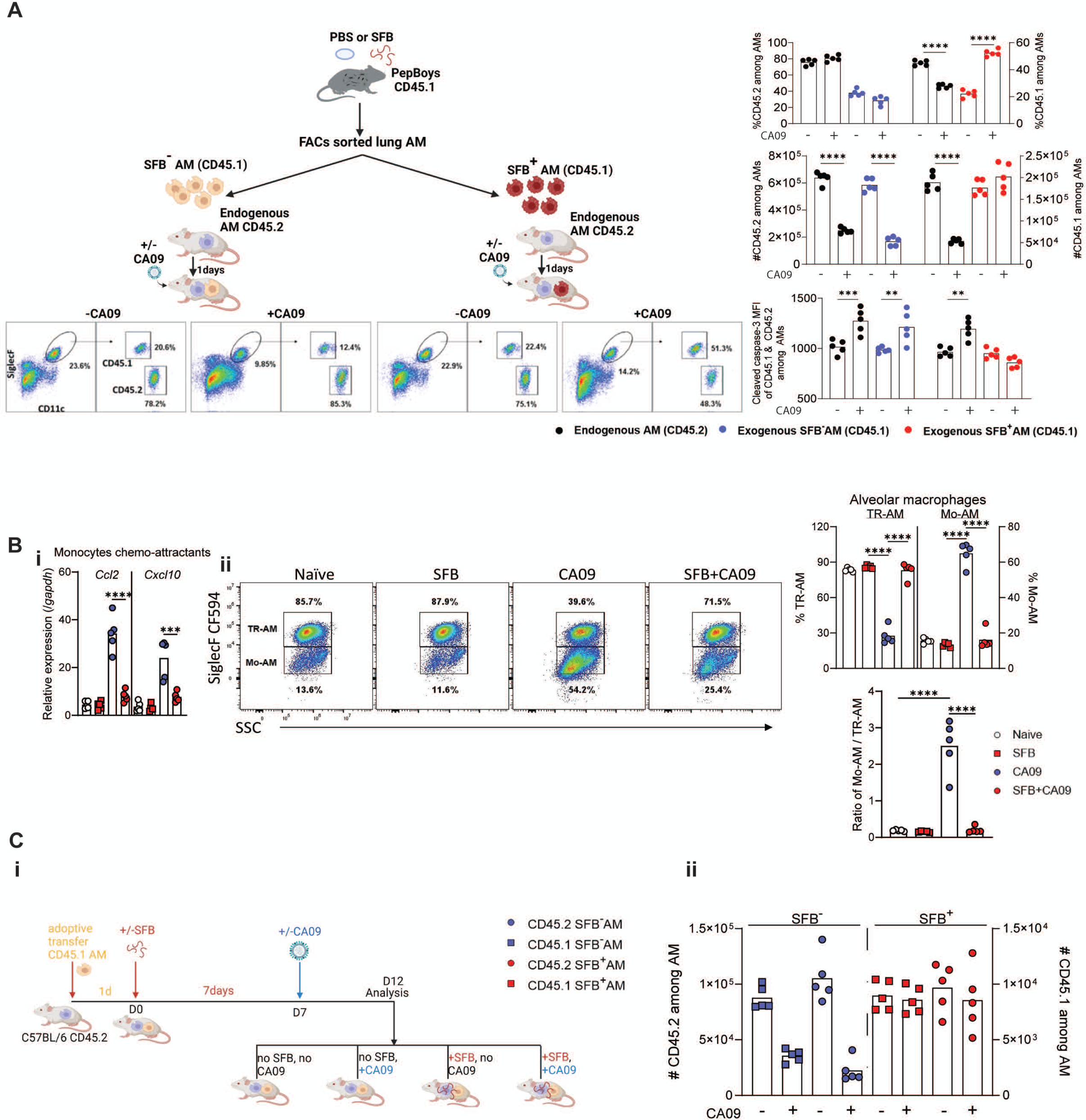
SFB colonization intrinsically altered AM to withstand IAV-induced depletion. **(A)** As schematized, SFB^-^ CD45.1 mice were administered with SFB or PBS. Seven days later, AM were isolated by FACs-sorting, and transferred to SFB-CD45.2 mice. One day post-transfer mice were inoculated with CA09. Four days later, lungs were harvested, and AM analyzed by FACs. Representative FACs plots, cell frequencies, cell numbers, and cleaved caspase-3 level are shown. **(B)** SFB^-^ mice were untreated (naïve) or administrated SFB. Seven days later, mice were inoculated (or not) with CA09. Mice were euthanized 4.5-days later. **(i)** qRT-PCR analysis of monocytes chemo-attractants *Ccl2* and *Cxcl10* from lung tissues. **(ii)** Representative FACs plots, cell frequencies, and ratio of tissue residents AM (TR-AM), and monocyte-derived AM (Mo-AM). **(C) (i)** As schematized, AM isolated from CD45.1 mice were transferred to CD45.2 mice. One day post-transferred, mice were administered with SFB or PBS for seven days then mice were inoculated with CA09. For day later, lungs were harvested, and AM analyzed by FACs. **(ii)** Cell numbers are shown. All experiment n = 5 mice per group. Data is representative of two independent experiments (A-B), yielding an identical pattern of results. Statistical test: One-way ANOVA. **p<0.01, ***p<0.001, ****p<0.0001, ns not significant.

The ability of transplanted SFB^+^ AM to withstand IAV-induced depletion could potentially have reflected that, in the donor mice, SFB colonization had phenotypically transformed resident AM and/or led to these cells being replaced by newly recruited monocytic precursors, which then became AM. That SFB^+^ mice did not exhibit a basal or IAV-induced elevation of monocytes chemo-attractants or monocyte-derived AM, both of which were readily evident in IAV-infected SFB^-^ mice, argued against the “AM replacement” mechanism (**Figure 5B**). Furthermore, administering SFB to mice after AM transplant led to similar persistence of both CD45.2 (endogenous) and CD45.1 (exogenous) cells following IAV infection (**Figure 5C**), thus strongly indicating that SFB colonization changed the phenotype of lung resident AM.

### SFB^+^ AM exhibited inflammatory anergy and enhanced C1qa-mediated antiviral function

Differences between SFB^+^ and SFB^-^ AM were characterized *ex vivo* as schematized (**Figure 6A**). Profiling of basal gene expression (i.e. without IAV exposure) by RNA-seq found that only 24 of the 15,510 genes expressed by AM were differentially expressed between SFB^-^ and SFB^+^ AM (**Figure 6B**), 18 of which were both confirmed by PCR and biologically replicated in a distinct mouse cohort **(Figure S6A).** Six of the genes upregulated in SFB^+^ AM are associated with an M2 macrophage phenotype. Yet, quantitation of surface expression markers previously used to distinguish M1 vs M2 phenotypes ^25^ did not reveal differences between SFB**^+^** and SFB^-^ AM **(Figure S6B)**. The genes most prominently up- and down-regulated, respectively, in SFB^+^ AM were Tsc22d3 and Notch-4, which, respectively, function in suppressing and activating pro-inflammatory gene expression ^26,27^, suggesting such signaling might be dampened in SFB^+^ AM. Indeed, we found that SFB^+^ AM lacked the robust UV-irradiated IAV-induced activation of pro-inflammatory gene expression exhibited by SFB^-^ AM (**Figure 6C and Figure S7C)**. Such inflammatory anergy was partially recapitulated by antibody-mediated neutralization of Notch-4 suggesting that the SFB-induced reduction in AM Notch-4 expression in SFB^+^ AM contributed to this *ex vivo* phenotype and, moreover, the reduced inflammation observed in IAV-infected SFB^+^ mice. As noted above, in SFB^-^ mice, IAV infection resulted in depletion of AM, which are then replaced by newly recruited monocytes, some of which will acquire AM cell surface markers. Such “replacement AM” lacked any hint of inflammatory anergy. Rather, they exhibited robust inflammatory responses to IAV, similar to those of the AM they replaced (Figure S6H).

**Figure 6.**
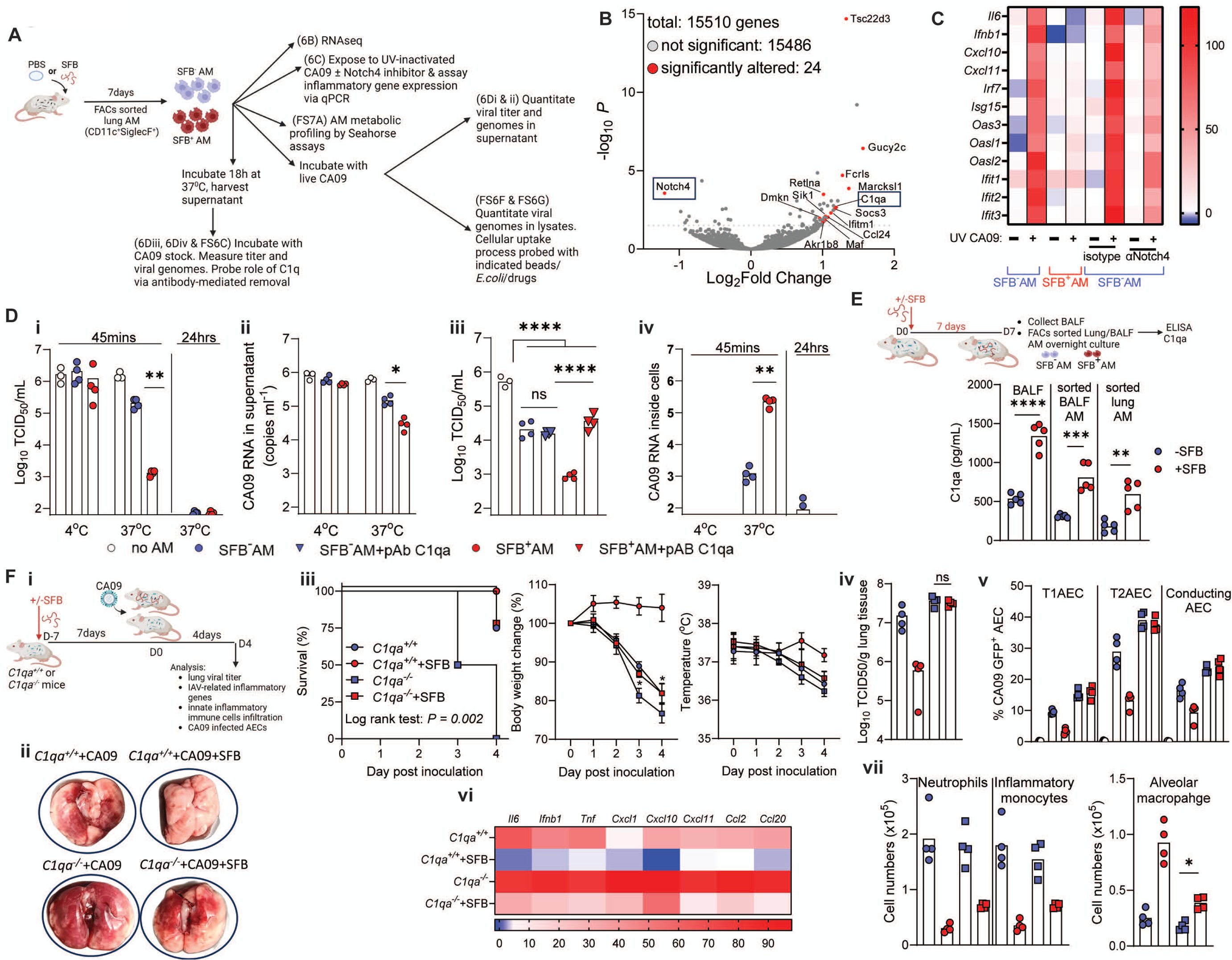
SFB^+^ AM exhibited inflammatory anergy and enhanced antiviral function *ex vivo*. **(A)** Schematic design for *ex vivo* studies of AM. **(B)** Comparison of basal SFB^+^ and SFB^-^ AM gene expression by RNA-seq, shown as a volcano plot. **(C)** SFB^+^ and SFB^-^ AM were exposed to UV-CA09 with or without Notch4 neutralizing, or isotype control antibodies. RNA was harvested 24 hours later, and gene expression assayed by RT-qPCR shown as a heatmap. **(D)** AM and CA09 stocks were incubated at 37°C, or indicated temperature, for 45 minutes and supernatants were collected. In addition, some cells had new media added and were incubated for another 24 hours. **(i)** Vius infection titers in supernatant at 45 minutes and 24 hours were assayed. **(ii)** Supernatants were collected after 45 minutes incubated with CA09 and viral genomes quantitated by RT-qPCR. **(iii)** Cell-free supernatant of 15 hours AM cultures were incubated with live CA09 stocks, with or without C1qa-neutralization or isotype antibody for 45 minutes, at which point virus infection titers were assayed. **(iv)** Cell lysates were generated upon CA09 removal or 24 hours later and viral genomes quantitated by RT-qPCR. **(E)** As schematized, mice were administered with SFB. Seven days later, bronchoalveolar lavage fluid (BALF) and lung were harvested, and AM were isolated via FAC-sorting. C1qa secretions were assayed by ELISA. **(F) (i)** 15 weeks old *C1qa^+/+^* and *C1qa^-/-^* SFB^-^ mice were untreated (naïve) or administrated SFB and then, 7 days later, inoculated with CA09. Mice were euthanized and lungs were harvested 4.5-days post CA09 inoculation. **(ii)** Images of lungs. **(iii)** Survival, body weight change, temperature. **(iv)** Lung viral titer. **(v)** Frequencies CA09 level in T1ACEs, T2AECs, and Conducting AECs were assayed by FACs. **(vi)** Gene expression assayed by RT-qPCR shown as a heatmap. **(vii)** Cell numbers of neutrophils, inflammatory monocytes and AM assayed by FACs. Statistical analyses of survival, body weight change, and temperature were conducted between *C1qa^-/-^* and *Cq1a^-/-^*+SFB group. All experiment n = 4-5 mice per group. Data is representative of two independent experiments, yielding an identical pattern of results. Statistical analysis: One-way ANOVA. *p<0.05, **p<0.01, ****p<0.0001, ns not significant.

The small panel of genes upregulated in SFB^+^ AM also included some known to function in antiviral resistance, including complement C1qa and Ifitm1, thus suggesting that SFB^+^ AM were not anergic to IAV per se but, rather, might have better virus neutralizing capacity. We examined this possibility by incubating CA09 stocks with AM for 45 min at which time AM were removed by centrifugation and supernatant infectious titers and IAV genomes quantified. SFB^-^ AM lowered the infectious titers of the CA09 stocks to which they were exposed, at 37°C but not 4°C, but the extent of reduction was 150-fold greater for SFB^+^ AM (**Figure 6Di**). This greater reduction in titer was accompanied by only 5-fold lower levels of PCR-quantifiable IAV genomes in supernatants of SFB^+^ AM (**Figure 6Dii**). This led us to hypothesize enhanced neutralization of IAV by complement and/or other products secreted by SFB^+^ AM. Indeed, we found that, even when SFB^+^ AM were not exposed to virus, their supernatants from 18h cultures had greater capacity than those from SFB^-^ AM to lower CA09 infectious titers (**Figure 6Diii**). Such enhanced ability of SFB^+^ AM supernatants to lower IAV titers was fully reversed by antibody-mediated removal of C1qa and, moreover, was not observed in C1qa^-/-^ SFB^+^ AM **(Figure S6C)**. These results accord with recent observations that C1qa can directly bind IAV and reduce its infectivity ^28^. In contrast to AM themselves, which when incubated with CA09 stocks lowered both levels of infectious virus (i.e. titer) and levels of viral RNA, AM supernatants did not impact levels of viral RNA (**Figure S6E**) suggesting the enhanced antiviral capacity of SFB^+^ AM might also involve increased viral absorption and/or uptake, possibly as a result of viral particles being more decorated with C1qa ^29^. In accord with this notion, reductions in IAV genomes in supernatants of AM/CA09 mixtures were paralleled by increased IAV genomes in AM lysates (**Figure 6Div**). The relative increase in IAV genomes in SFB^+^, compared to SFB^-^, AM lysates was maintained amidst chlorpromazine, but not latrunculin treatment, neither of which impacted cell viability, suggesting it might reflect enhanced phagocytosis uptake by SFB^+^ AM (**Figure S6G**). In contrast to our hypothesis, increased uptake of CA09 was evident in C1qa^-/-^ SFB^+^ AM (Figure S6D) suggesting SFB^+^ AM may be more phagocytic in general. In accord with this notion, SFB^+^ AM exhibited modestly greater uptake of IgG-coated fluorescent beads *ex vivo* and bioparticles *in vivo* **(Figure S6F**). Exploring the fate of CA09 internalized by AM during the 45 min exposure period found that, 24 h following media replacement, AM supernatants had few infectious virions while lysates contained few detectable IAV genomes (**Figure 6Div**) thus suggesting that AM were not permissive for CA09 replication, but rather constituted a dead end for the virus. Collectively, these results indicate that elevated C1qa expression and enhanced phagocytosis drove the increased antiviral capacity of SFB^+^ AM.

Measure of C1qa protein by ELISA confirmed that SFB colonization increased expression of C1qa in AM isolated from whole lung and bronchoalveolar lavage fluid (BALF) and, moreover in whole BALF (**Figure 6E**). Furthermore, experiments in *C1qa^-/-^* mice indicated that C1qa played a central role in SFB’s suppression of IAV in vivo. Specifically, in *C1qa^-/-^*mice, SFB colonization did not lower viral titers or protect lung epithelial cells from IAV infection following CA09 inoculation (Figure 6F). Concomitantly, SFB’s ability to ameliorate disease symptoms, including weight loss and hypothermia was markedly reduced in *C1qa^-/-^* mice relative to *C1qa^+/+^*mice. Nonetheless, colonization of *C1qa^-/-^* mice with SFB still conferred modest but statistically significant reductions in weight loss and mortality. Moreover, SFB colonization of *C1qa^-/-^* suppressed CA09-induced pro-inflammatory gene expression, gross lung pathology, and infiltration of inflammatory cells into the lungs. The reduced but not eliminated ability of SFB to suppress symptoms of IAV infection in *C1qa^-/-^*mice associated with SFB colonization partially protecting AM against IAV-induced depletion. Thus, C1qa was germane to SFB’s ability to impede IAV infection and its consequences but nonetheless SFB ameliorated IAV-induced lung inflammation irrespective of C1qa and viral titer.

### SFB-induced reprogramming of AM protects mice against IAV infection

The starkly different transcriptional responses of SFB^-^ and SFB^+^ AM led us to investigate the general metabolic states of these cells via Seahorse analysis. The basal metabolic states of SFB^-^ and SFB^+^ AM differed only modestly with SFB^+^ AM exhibiting a small reduction in glycolysis (Figure S7A). In contrast, inoculating SFB^-^ mice with CA09 led to stark increases in AM ATP production and glycolysis, which were wholly absent in SFB^+^ mice thus paralleling above-described changes in inflammatory parameters in UV-IAV induced changes in AM proinflammatory gene expression observed *ex vivo*. Lastly, we examined the extent to which SFB-induced reprogramming of AM contributed to SFB’s protection against IAV infection. Specifically, we performed an additional series of AM transplant experiments designed to measure impacts on IAV levels and clinical-type parameters. Consistent with the notion that AM, prior to their depletion, protect against IAV infection ^24^, we observed that administration of SFB^-^ AM to a host replete with endogenous AM resulted in a modest reduction in IAV titers, gross lung damage, hypothermia, weight loss, and death following CA09 inoculation (**Figure 7A**). Administration of SFB**^+^** AM provided markedly greater protection against this IAV challenge, approximating the level conferred by administration of SFB itself, as assessed by all parameters measured. Analogous patterns of results were seen following transfer of SFB^+^ and SFB^-^ AM into mice deficient in endogenous AM, achieved via use of *Csf2^-/-^* mice or chlodronate administration (**Figure 7B and 7C**). Analogous to results obtained in *C1qa^-/-^* mice, transplant of C1qa^-/-^ SFB**^+^** AM into SFB^-^ mice did not reduce viral titers or protect lung cells from IAV infection following CA09 inoculation but nonetheless reduced indices of inflammation and conferred a modest clinical-type benefit **(Figure S7B)**. *Ex vivo* blockade of Notch-4 on SFB^-^ AM prior to transplanting them also enhanced their ability to withstand IAV-induced depletion, protect EC from IAV infection, and ameliorate CA09-induced disease, but did not impact level of viral genomes in the lung **(Figure S7C)**, suggesting the ability of transplanted SFB^+^ AM to do so was not mediated solely by dampened inflammatory signaling but also required such AM have enhanced antiviral function. Thus, SFB colonization of the gut stably and broadly altered the AM phenotype enabling these cells to better protect their hosts from lethal RVI.

**Figure 7.**
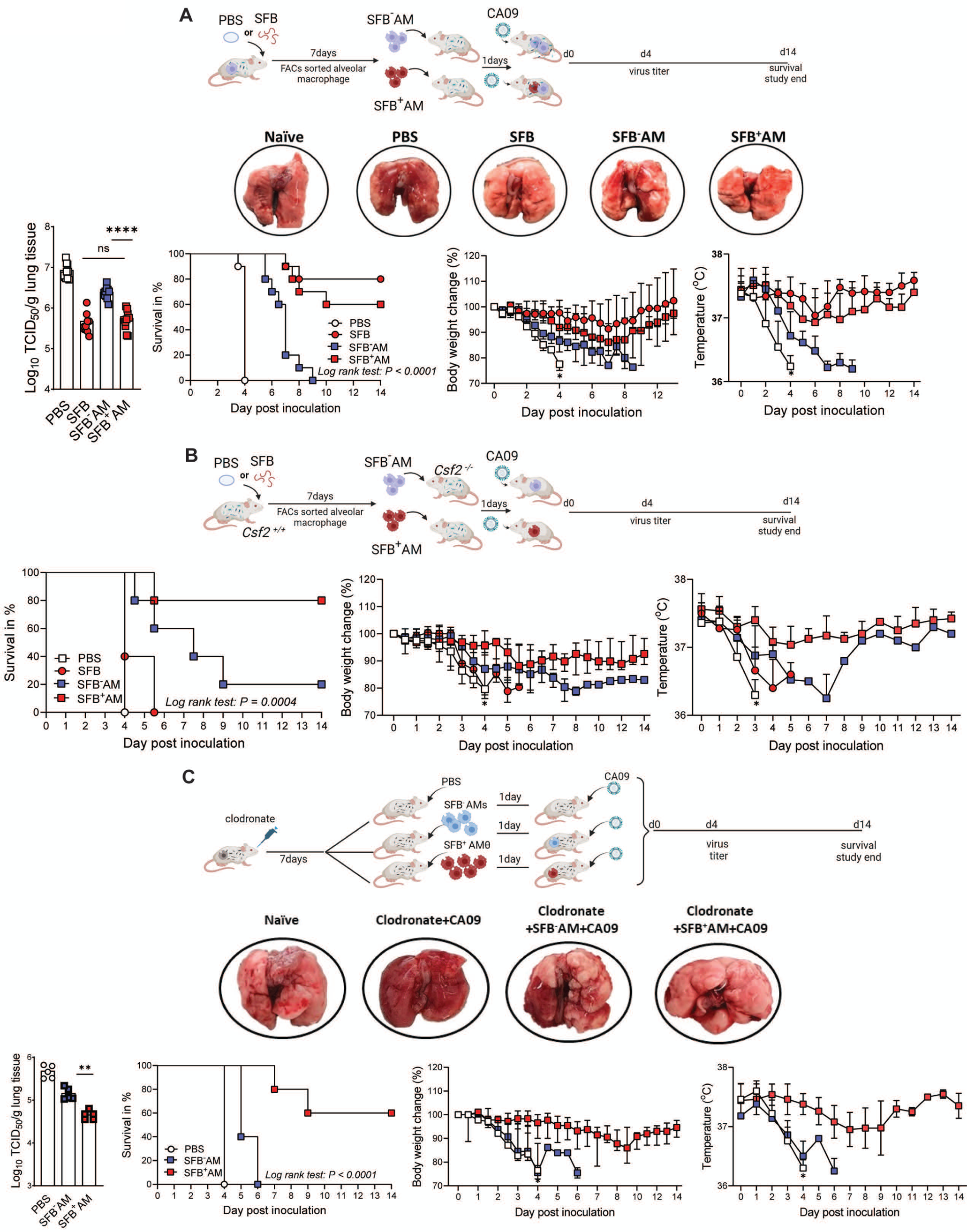
Adoptive transfer of SFB^+^ AM enabled recipients to better manage IAV infection. **(A)** As schematized, SFB^-^ mice were administered SFB or PBS. Seven days later, AMs were isolated by FACs-sorting and transferred to SFB^-^ mice. Mice were administered 5×10^5^ cells per mouse. One day post adoptive transfer, mice were inoculated with a lethal dose of CA09. Mice were euthanized 4.5 days post-inoculation for viral titer and gross lung assessment or monitored for survival, body weight, and temperature. Statistical analyses of survival, body weight change, and temperature were conducted using the “PBS” group as the control group. **(B)** As schematized, AMs from SFB^-^ *csf2^+/+^* mice were administered SFB or PBS. Seven days later, AMs were isolated by FACs-sorting and transferred to SFB^-^ *csf2^-/-^* mice. One day post-transfer, mice were inoculated with CA09. Mice were monitored for 14 days for clinical symptoms including survival rate, body weight, and temperature. **(C)** SFB^-^ mice were administered clodronate liposomes to remove endogenous AM populations. Seven days later after initiation of depletion, mice were administered 5×10^5^ AM cells harvested from the SFB^-^ or SFB^+^ mice. One day post transfer, mice were inoculated with CA09. Mice were euthanized on day four to assess lung viral titer and gross lung assessment or monitored for survival, body weight, and temperature. All experiments have at least n = 5 mice per group. Data is representative of two independent experiments (A), yielding an identical pattern of results. Statistical analyses of survival, bodyweight change, and temperature of (A-C) were conducted by comparing “PBS” group to all other groups. Statistical analysis; One-way ANOVA: viral titers. Survival: Log-rank Mantel-Cox test. Body weight, temperature: Two-way ANOVA. *p<0.05, **p<0.01, ****p<0.0001, ns not significant.

## DISCUSSION

Positioned in the lung lumen to enable their guarding the surfaces of O_2_ and CO_2_ exchange, AM are amongst the first cells to encounter inhaled threats including respiratory viruses. We show herein that how AM respond to respiratory viruses is dramatically altered by the composition of the intestinal microbiota. Specifically, we report that colonization of the intestine by SFB, naturally acquired, or exogenously administered, markedly attenuates IAV-induced AM pro-inflammatory gene expression while increasing ability of AM to disable IAV. This seemingly paradoxical phenotypic differences between SFB^-^ and SFB^+^ AM is reminiscent of the comparison of blood monocytes and intestinal macrophages wherein the latter are more phagocytic but yet exhibit inflammatory anergy ^30^. An important aspect of SFB-induced reprogramming of AM is that it resulted in resistance of RVI-induced depletion and thus sustained protection of lung epithelial cells. Collectively, the consequences of SFB-induced AM reprogramming resulted in lower viral titers, reduced inflammation, and reduced disease severity.

While the general concept that microbiota composition in total can influence immune phenotype and, proneness to infection is established ^31,32^, we were surprised that the presence of a single common commensal bacterial species, amidst the thousands of different microbial species that variably inhabit the mouse gut, had such strong impacts in RVI models and, furthermore, that such impacts were largely attributable to reprogramming of AM. The extent to which SFB is present in humans is not well defined but it has been reported to be frequently present in children in some regions ^33^ and was unequivocally shown to be present in some adult individuals ^34^. Independent of a specific role for SFB, however, our findings speak strongly to the potential of gut microbiota to influence proneness to RVI, especially via programming of AM. Both AM pro-inflammatory gene expression and AM depletion are known to correlate with RVI severity in humans but determinants of inter-individual heterogeneity in AM responses have not been defined ^33^. We consider it highly unlikely that SFB is the only gut microbe capable of impacting AM phenotype and, consequently, proneness to RVI. Rather, we posit that AM are a key component of the gut-lung axis wherein an individual’s microbiota in its entirety is a determinant of disease outcome following exposure to an array of respiratory viruses.

The observation that SFB**^+^** AM maintained an IAV-resistant phenotype following transplant to SFB^-^ mice is consistent with observations that AM have stable cell-type specific patterns of gene expression and methylation even when cultured extensively *ex vivo* ^35^. Moreover, it accords with recently increasing appreciation that AM can be “trained” to display long-lasting phenotypic alterations that impact their ability to manage challenges. Yet the consequences and mechanisms of previously observed AM training seems quite distinct from that reported herein. For example, administration of live, but not heat-inactivated BCG vaccine resulted in systemic low-grade infection and inflammation that altered gut microbiota composition and increased gut permeability, potentiating AM pro-inflammatory gene expression ^36^. In addition, compared to BCG, SFB colonization had a somewhat opposite outcome on AM function in that it suppressed rather than promoted pro-inflammatory gene expression. In further contrast to BCG, phenotypic impacts of SFB were maintained in gnotobiotic mice, arguing against broad changes in microbiota composition playing a central role. Another starkly different example of AM training is the recent observation that IAV infection increases ability of AM to suppress tumors ^37^. Such AM training is mediated by a large-scale depletion of AM, which are then replaced by cells newly recruited from bone marrow that, even after establishing residency in the lung, exhibit stark global differences in gene expression profiles. In contrast, SFB colonization phenotypically transformed resident AM without a detectable influx of monocyte precursors. Moreover, gene expression profiling of SFB^-^ and SFB^+^ AM revealed only modest differences, likely reflecting that both SFB^-^ and SFB^+^ AM had been generated by AM self-renewal processes that normally maintain AM homeostasis in non-traumatic states.

Despite being modest, SFB-induced changes in basal AM gene expression contributed mightily to the enhanced ability of SFB-colonized mice to manage IAB infection. Most notably, use of *C1qa^-/-^* mice, and AM transferred therefrom, indicated that SFB-induced increases in C1qa expression were pivotal for lowered viral titers. The comparable genetically-modified mice to similarly examine the necessity of genes that mediated SFB’s suppression of AM inflammatory signaling are not yet in hand. However, a converse approach, namely use of a Notch-4 neutralizing antibody, argued that the SFB-induced reduction in Notch-4 expression was germane to SFB-induced AM inflammatory anergy and that this aspect of the SFB^+^ AM phenotype also contributed to protection against IAV. In addition to directly disabling IAV, AM can also protect lung EC from IAV infection via suppressing EC CysLT1 signaling, which might mediate IAV entry into these cells ^24^. Thus, persistence of AM in SFB^+^ mice, by itself, likely also contributed to SFB’s protection against IAV. That mechanisms driving RVI-induced AM depletion are not clear, makes determining how SFB colonization prevents such depletion challenging. Indeed, deciphering the interrelationships between reduced viral titers, inflammatory anergy, and increased AM survival, is a complex endeavor in which some events are likely both a cause and consequence of each other. For instance, collectively our results indicate that AM persistence enabled them to suppress IAV, but yet such suppression of IAV likely contributed to increased AM survival. Conversely, we envisage that SFB-induced AM inflammatory anergy promoted AM persistence, which in turn enabled sustained protection of epithelial cells from IAV infection. Such protection of epithelial cells from IAV infection would reduce the viral burden, which likely contributes to both inflammation and AM death. Hence, defining the relative contributions of enhanced antiviral and suppressed inflammation is a complex undertaking. Nonetheless, our current working hypothesis is that elevated C1qa expression plays a leading role, and inflammatory anergy plays a key supporting role, in mediating SFB’s protection against RVI.

Less clear is the nature of the signal that is generated by SFB colonization of the luminal surface of the intestine that drives AM reprogramming. We speculate that SFB produces, or induces host production of, a metabolite(s) that reach the lung and act upon AM, changing both their basal gene expression and how they respond to challenge, perhaps by altering DNA methylation. Additionally/alternatively, it is possible that gut to brain and/or brain to lung neural signaling is involved. We envisage that deciphering how SFB impacts lung AM may yield strategies to mitigate disease burden cause by RVI. Indeed, in addition to mitigating primary RVI, AM persistence may help prevent secondary consequences of RVI. Many of the severe consequences of IAV infection in humans result from secondary bacterial infections, to which the lung is more permissive by loss of IAV-induced AM depletion ^38^. Such impaired antibacterial immunity can persist for prolonged periods perhaps reflecting that the bone-marrow derived replacement AM are not quickly optimized for mucosal host defense. Thus, microbiota mediated AM programming may not only influence severity of acute RVI but may also be a long term post-RVI health determinant.

## Acknowledgments

We thank Tim Denning and Ifor Williams for insightful guidance. We thank RS Baric for SARS-CoV-2 MA10 virus isolate and Georgia State University’s High Containment Core and Division of Animal Resources for support.

## Author contributions

Conceptualization: RKP, ATG. Investigation: VLN, CML, HK, KS. Resources: RKP, ATG. Visualization: VLN, CML, KS, RKP, ATG. Validation: RKP, ATG. Funding acquisition: RKP. Project administration: RKP, ATG. Supervision: RKP, ATG, MK. Writing – original draft: ATG. Writing – review & editing: VLN, CML, KS, RKP.

## Data and materials availability

All data are available in the main text or the supplementary materials.

## STAR METHODS

### RESOURCE AVAILABILITY

#### Lead contact

Further information and requests for resources and reagents should be directed to and will be fulfilled by the lead contact, Andrew Gewirtz.

#### Materials availability

This study did not generate new unique reagents.

#### Data and code availability

- The raw sequencing data has been deposited at GenBank with access number: PRJEB71689, and PRJEB71449, and all other data used for this study will be shared upon reasonable request from the lead contact.
- This paper does not report original codes.
- Any additional information required to reanalyze the data reported in this paper is available from the lead contact upon reasonable request.

## EXPERIMENTAL MODEL DETAILS

### Virus strains and growth conditions

All viruses and cells used in this study are listed in the key resources table. Madin-Darby Canine Kidney (MDCK; ATCC CCL-34), HEp-2 (ATCC CCL-23), and VeroE6 TMPRSS2 (BPS Bioscience #78081) cells were cultured in Dulbecco’s Modified Eagle’s Medium (DMEM) supplemented with 7.5% heat-inactivated fetal bovine serum (FBS) at 37°C and 5% CO_2_. All cell lines used in this study were authenticated and checked for *Mycoplasma* and microbial contamination. Recombinant Influenza A virus was recovered based on original reports for laboratory adapted IAV ^39^. A/CA/07/2009 (H1N1) was grown on MDCK cells using serum-free DMEM supplemented with 0.5% Trypsin. Recombinant RSV strain A2 with line19F reporter gene recRSV-mKate was rescued and amplified as described previously ^40^, and the SARS-CoV-2 MA10 strain ^41^ was passaged in VeroE6 TMPRSS2 cells. All viruses used were clarified, aliquoted, and stored at -80°C. Virus stock titers were determined through TCID_50_-titration.

### Complex microbiota and mono-associated SFB transplantation

Fecal samples were collected from donor mice, suspended in 20% glycerol PBS solution at 40 mg/mL concentration, passed through a 40 µm filter, and stored at -80C. Frozen fecal suspensions were orally administered to recipient mice at 200 µl per mouse.

### Virus infections

All viruses and mice strains used in this study are listed in the key resources table. The Institutional Animal Care and Use Committee of Georgia State University approved animal studies. All mice, unless otherwise stated, were purchased from The Jackson Laboratory. The Jackson Laboratory does not perform testing for segmented filamentous bacteria (SFB) in mice. Nevertheless, all mice obtained from this vendor were tested and confirmed to be SFB-negative upon their arrival at GSU. The following mice were purchased from Taconic Biosciences: C57BL/6 (B6 WT) Excluded Flora (EF) mice and Specific Pathogen-Free (SFP) mice, which we verified were SFB+ and SFB-, respectively. Unless otherwise stated, mice were 5 weeks of age at the time of infection. Upon arrival, mice were acclimated for five days and then randomly assigned to study groups and housed under ABSL-2 or ABSL-3 conditions for infections with A/CA/07/2009 (H1N1) recombinant, recRSV-mKate, or SARS-CoV-2 MA10 strain, respectively. Seven days prior to virus infection, mice were subjected to microbiota manipulation (administration of feces from SPF or SFB-monoassociated mice). Bodyweight was determined twice daily, and body temperature determined once daily rectally. For infections, animals were anesthetized with isoflurane [rec A/CA/07/2009 (H1N1) (CA09)] or ketamine/xylazine (recRSV-mKate and MA-SARS-CoV-2), followed by intranasal inoculation with 2.5×10^2^-2×10^4^ TCID_50_ units/animal of GFP-expressing reporter virus version of A/CA/07/2009 (H1N1) (CA09), 5.5×10^5^ of recRSV-mKate, or 4.2×10^5^ of MA-SARS-CoV-2. In all cases, virus inoculum was diluted in 50 µl and administered in amounts of 25 µl per nare, followed by treatment with 4’-FlU at specified dose levels and starting time points through oral gavage. Each individual study contained animals receiving equal volumes of vehicle through oral gavage. Animals were euthanized and organs harvested at predefined time points or when animals reached humane study endpoints. At study end point, lung tissue was harvested for virus titration or histopathology.

## METHODS DETAILS

### PCR primers, FACs antibodies, neutralization antibodies and other reagents

All reagents used in these manuscripts can be seen in the key resources table.

### Histopathologic analysis

Lungs were perfused using 10% neutral-buffered formalin, dissected, and fixed. Formalin-fixed lungs were transferred to 70% EtOH, embedded in paraffin, sectioned, stained, and scored by a board-certified veterinary pathologist (KS), who was blinded to the study groups. For each parameter, the most severe lesion in the section was scored. Pleuritis, bronchiolitis, and alveolitis scoring was based on distribution: 1 = focal, 2 = multifocal, 3 = multifocal to coalescing, 4 = diffuse. For interstitial pneumonia (IP): 1 = alveolar septa expanded by 1 leukocyte thickness, 2 = 2-leukocytes thick, 3 = 3-leukocytes thick, 4 = 4-leukocytes thick. For vasculitis: 1 = vessel wall infiltrated by leukocytes, 2 = infiltration and separation of myocytes, 3 = same changes as in 2 with fibrinoid change. For perivascular cuffing (PVC): 1 = vessels surrounded by 1 leukocyte layer, 2 = 2 to 5-leukocytes thick, 3 = 6 to 10-leukocytes thick, 4 = more than 10-leukocytes thick.

### Virus titration

Virus samples were serially diluted (10-fold starting at 1:10 initial dilution) in serum-free DMEM (recRSV-mKate and MA-SARS-CoV-2) and supplemented with 0.5% Trypsin (Gibco, recA/California/07/2009). Serial dilutions were added to cells seeded in 96-well plate at 1×10^4^ cells per well, 24 hours before infection. Infected plates were incubated for 72 hours at 37°C with 5% CO_2_ and scoring of wells based on reporter activity/syncytia.

### Virus titration from tissue samples

Organs were weighed and homogenized in 300 µl PBS using a bead, set to 3 cycles of 30 seconds each at 4°C, separated by 30-second rest periods. Homogenates were cleared (10 minutes at 20,000 × g and 4°C), and cleared supernatants were stored at -80°C until virus titration. Viral titers were expressed as TCID_50_ units per gram input tissue.

### Neutralization of IL-17 and IL-22

IL-17 (100 µg) and IL-22 (150 µg) neutralizing monoclonal antibody were administered intraperitoneally before microbiota transplantation and virus inoculation thereafter every other day until mice were euthanized.

### Lung digestion and flow cytometry analysis/isolation of innate immune cells

Lungs were dissected from mice and finely minced with scissors before digestion (40 minutes, at 37°C) with warm RPMI containing 10% FBS, DNase, and 1mg/mL Collagenase type IV (Sigma); lungs were further homogenized via swishing with a 3-mL syringe every 10 minutes during digestion. Following digestion, immune cells were purified with 40%/80% isotonic Percol gradient centrifugation. For flow cytometry analysis, the prepared cells were blocked with 1 µg/million cells with 2.4G2 (anti-CD16/anti-CD32) in 100 µL PBS for 15 minutes at 4°C, followed by 2-times wash with PBS to remove residue of 2.4G2 and then incubated in conjugated mAbs CD45 BV605, MHCII BV650, CD11b APC-Cy7, CD11c BV786, Ly6C PerCp-Cy5.5, Ly6G AF700, CD64 BV421, CD24 FITC, CD117 GV510, SiglecF PE-CF594, Fcer1 APC. Intracellular staining for cleaved caspase-3 was done using FoxP3 staining buffer purchased from eBioscience. Multi-parameter analysis was performed on a CytoFlex (Beckman Coulter) and analyzed with FlowJo software (Tree Star). Cell sorting was done using Sony SH800Z cell sorter (Sony). Gating strategy for innate immune cells was based on Yu *et al*., ^42^ and gating strategy for airway epithelial cells was based on Cardani *et al*., ^24^. Immune cells and lung airway epithelial subtypes are defined in Table S1.

### RNA-Seq

RNA was isolated from lungs or FACs-sorted alveolar macrophages using RNeasy Plus Mini Kit from Qiagen. The prepared RNA samples were then sent to the Molecular Evolution Core of Georgia Institute of Technology for library preparation and sequencing on the NextSeq instrument with a high output 2×75bp run. FASTQC was used for sequencing reads quality screening. The RNAseq data were then analyzed using the Galaxy server.

### Adoptive transfer of alveolar macrophages

Lungs were harvested from CD45.1 mice and digested with Collagenase type IV as described above. Alveolar macrophages (defined at CD45^+^MHCII^+^CD11c^+^Siglect^+^) were stained and FACs-sorted. Small aliquot of isolated cells was used to determined purity. Unless stated otherwise, 5×10^5^ alveolar macrophages in 30 µl PBS were administered intranasally to recipient mice. One day post cells transfer, some mice were euthanized to determine transfer efficiency and survival of transferred cells; and some mice were inoculated with IAV. For survival experiments, mice were monitored twice daily for 14 days. For lung virus titers, mice were euthanized on day 4.5 post virus inoculation.

### *Ex vivo* inhibiting Notch-4 signaling in alveolar macrophages

Alveolar macrophages were FACS-sorted as described. After sorting, cells were treated with either isotype control antibody (Cat# BE0091, BioXCell) or an anti-Notch4 monoclonal antibody (20µg/mL) (Cat# BE0129, BioXCell), known to neutralize Notch-4 [27] for 30 minutes. Cells were washed twice and resuspended in PBS and intranasally transferred to mice.

#### *Ex vivo* alveolar macrophages culture, virucidal assay, and anti-body mediated removal of C1qa

Alveolar macrophages were FACS-sorted as described. After sorted cells were incubated for 18h at 37°C. Cells were washed twice with serum-free media and were incubated with live CA09 (MOI = 1) in serum-free DMEM supplemented with 0.5% Trypsin (flu media) for 45min.

In addition, to assess whether alveolar macrophages are the terminal destination for IAV, some cells had new flu media added and were incubated for another 24h. Viral titer and genomes were determined via TCID_50_ and PCR methods.

#### Antibody-mediated removal of C1qa from *ex vivo* alveolar macrophages’ supernatants

Cell-free supernatants of 18h AM cultures were treated with C1qa polyclonal antibodies or isotype for 2 hours at 4°C and protein A/G was added and centrifuged. Supernatant was harvested and incubated with CA09 stocks for 45mins, at which point IAV infection titers were assayed.

#### Alveolar macrophages phagocytosis assay

*Ex Vivo* alveolar AM bead uptake assay was performed with Phagocytosis assay kit (Cat#500290, Cayman chemical). Alveolar macrophages were FACS-sorted as described. After sorting, 1×10^5^ cells were seeded in each well of a 96-well plate and incubated for 18h at 37°C. Cells were washed twice, and IgG-FITC beads were added and incubated for 2 hours at 37°C. After incubation period, cells were centrifuged, and supernatants were removed and resuspended FACS buffer. Level of cell fluorescence, an indicator of bead uptake, was measured via flow cytometry. For *In vivo* AM bioparticle uptake, SFB^-^ and SFB^+^ mice were administered pHrodo^TM^ *E.coli* BioParticles^TM^ conjugated (Cat# P35360, Invitrogen). Three hours later, mice were euthanized and BALF were harvested, and AM were labeled. Level of cell fluorescence, an indicator of bioparticle uptake, was measured via flow cytometry.

### Quantitative real-time PCR

Total RNA from lungs or from FACs-sorted alveolar macrophages were isolated using the Qiagen RNeasy Mini Kit with on-column DNase digestion according to manufacturer’s protocol. cDNA was generated using the Superscript First Strand Synthesis kit for RT-PCR and random hexamer primers (Invitrogen). RT-qPCR was performed with SYBR Green using StepOnePlus PCR system (Applied Biosystem), and all genes expression was normalized to GAPDH.

### C1qa ELISA

BALF was collected from SFB^+^ and SFB^-^ mice. Alveolar macrophages (AM) were sorted using fluorescence-activated cell sorting (FACS) from lung and BALF of SFB^+^ and SFB^-^ mice. Sorted AM were incubated in RPMI + 10% FBS overnight at 37°C in a cell culture incubator. BALF and supernatants from cultured AM were coated onto 96-well ELISA plates and incubated overnight at 4°C. Plates were washed with PBST to remove unbound material. Blocking was performed with 3% bovine serum albumin (BSA) in PBS at room temperature for one hour. Polyclonal antibody against C1qa was diluted at 1:5000 and added to the plate and incubated for 2h at room temperature. Secondary antibodies (conjugated to horseradish peroxidase, HRP) were added and incubated at room temperature for 1 hour. Tetramethylbenzidine (TMB) substrate was added to detect HRP enzyme conjugate activity. Absorbance at 450 nm was measured to quantify the presence of C1qa. Mouse C1qa recombinant protein was serially diluted to construct a standard curve for determining the concentration of C1qa in the samples. Absorbance values of the samples were compared with the standard curve to determine the concentration of C1qa in the BALF and cultured AM supernatants.

### AM metabolic assay

The real-time metabolism of AM was assessed using the Seahorse XF Cell Mito Stress Test Kit or the Seahorse XF Glycolytic Stress Test Kit from Agilent Technologies, following the manufacturer’s instructions. AM were FACS-sorted from SFB^+^ and SFB^-^ mice. The sorted cells were then seeded into Seahorse cell culture plates at a density of 1×10^5^ cells per well and cultured overnight in complete RPMI1640 medium. On the following day, AM were incubated with Seahorse assay medium (Agilent Technologies) for one hour. For the glycolysis stress test, AM were supplemented with 2mM glutamine. For the mito stress test, AM were supplemented with 10mM glucose and 1mM pyruvate. All reagents provided in the kits were resuspended in assay medium, and the final concentrations in the culture wells were as follows: 1.5 μM oligomycin, 4 μM FCCP, and 0.5 μM rotenone/antimycin A for the mito stress test, and 10 mM glucose, 1 μM oligomycin, and 50 mM 2-deoxy-D-glucose (2-DG) for the glycolytic stress test. The oxygen consumption rate (OCR) and extracellular acidification rate (ECAR) were measured on a Seahorse XF analyzer. Non-glycolytic ECAR is calculated as the average ECAR values after 2-DG treatment. Basal glycolysis is calculated as the average ECAR value prior to oligomycin treatment minus non-glycolytic ECAR. Max glycolysis is calculated as the average ECAR values after oligomycin and before FCCP treatment. Glycolytic reserve is calculated as max glycolysis minus basal glycolysis. Basal respiration is calculated as the average OCR value prior to oligomycin treatment minus non-mitochondrial OCR. Max respiration is calculated as the average OCR value after FCCP and before rotenone/antimycin A addition.

### Experimental schematics and graphical abstract

All experimental schematics and graphical abstract created using BioRender. https://biorender.com/

### Quantification and statistical analysis

Results were expressed as mean ± SEM. All data was plotted in GraphPad Prism version 10. Statistical significance was assessed by One-way ANOVA, Student’s t test, and Two-way ANOVA. Differences between experimental groups were considered significant if *p<0.05, **p<0.01, ***p<0.001, ****p<0.0001.

---

**Figure S1 (related F1).**
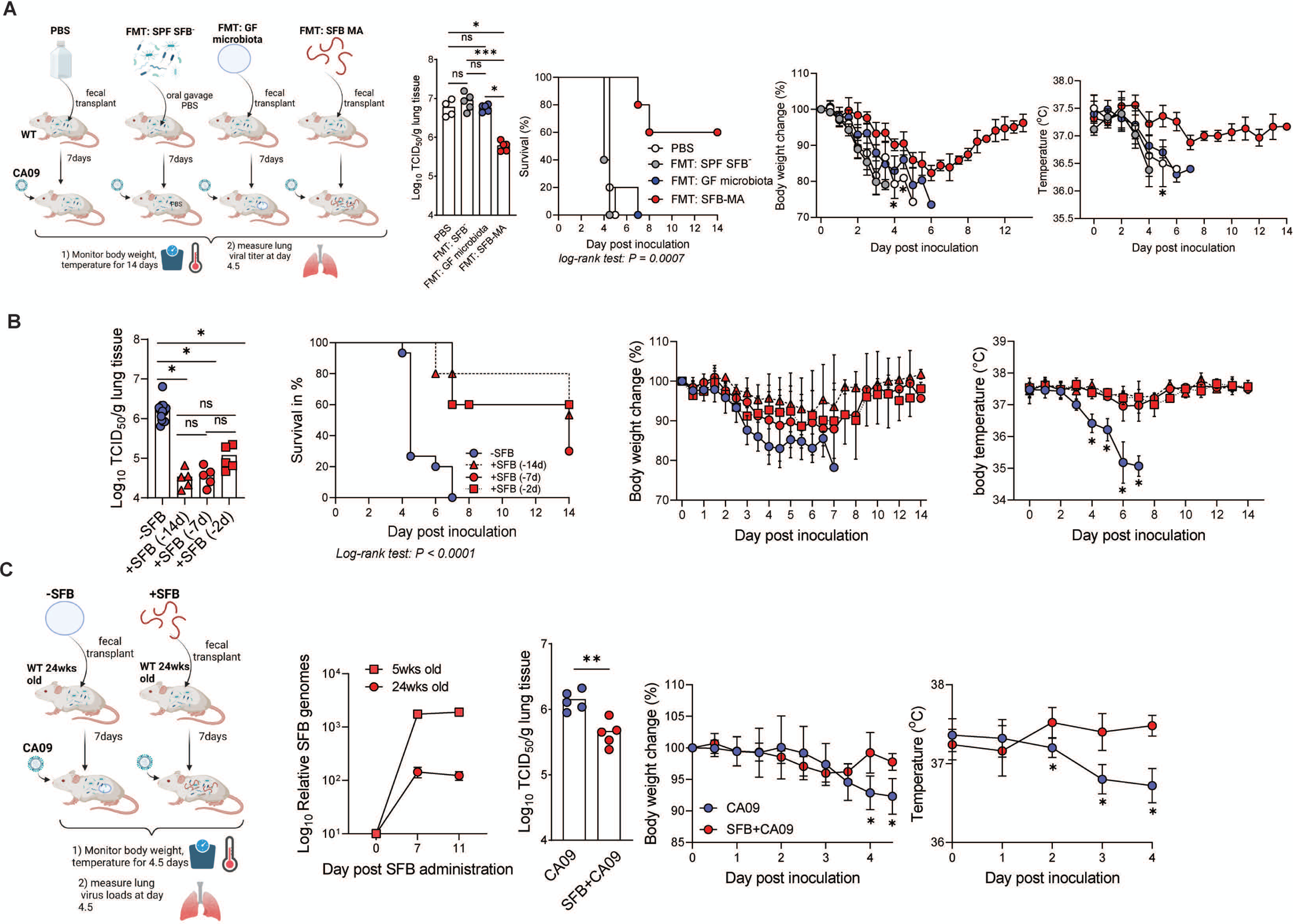

**Figure S2 (related F3).**
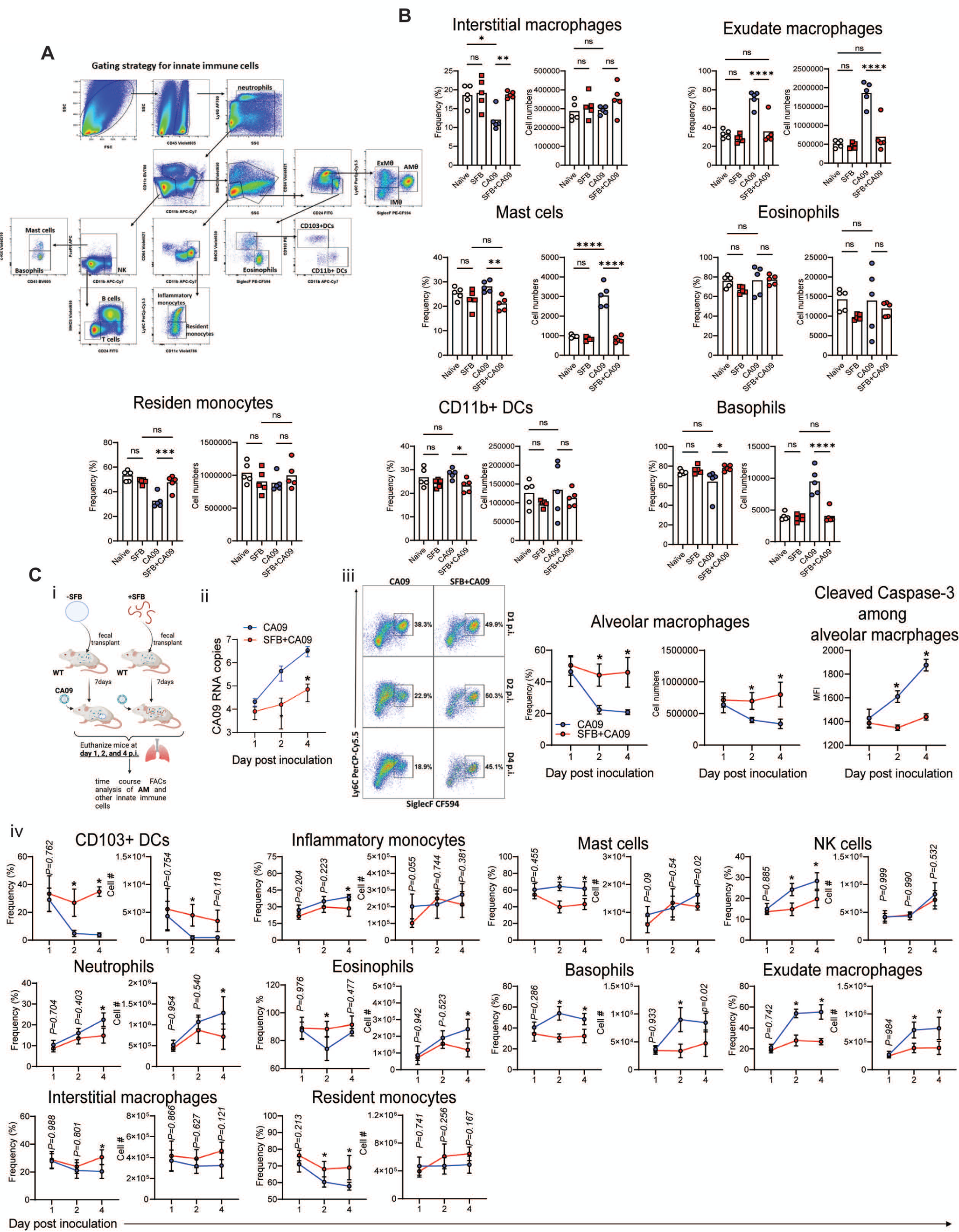

**Figure S3 (related F4).**
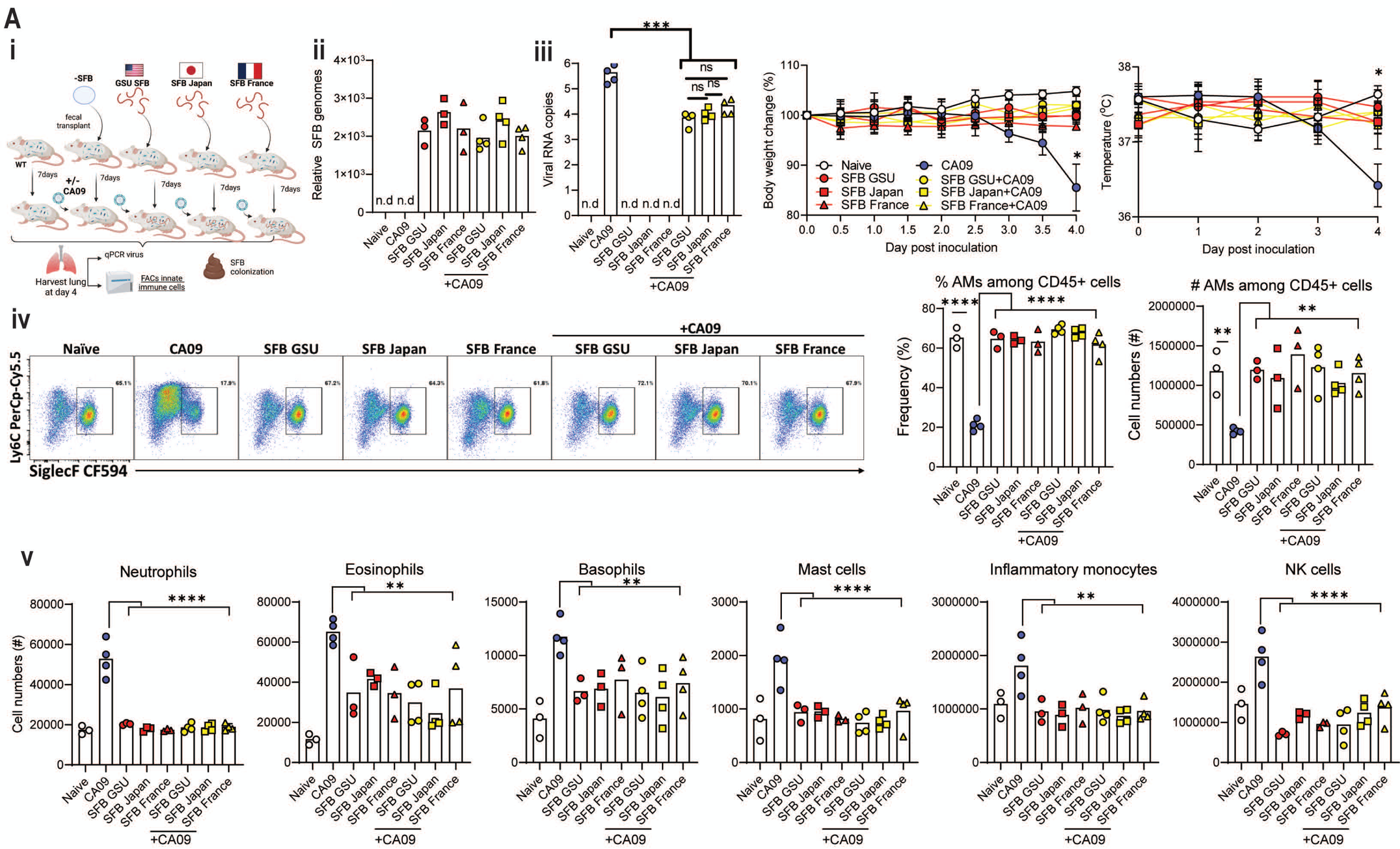

**Figure S4 (related F4).**
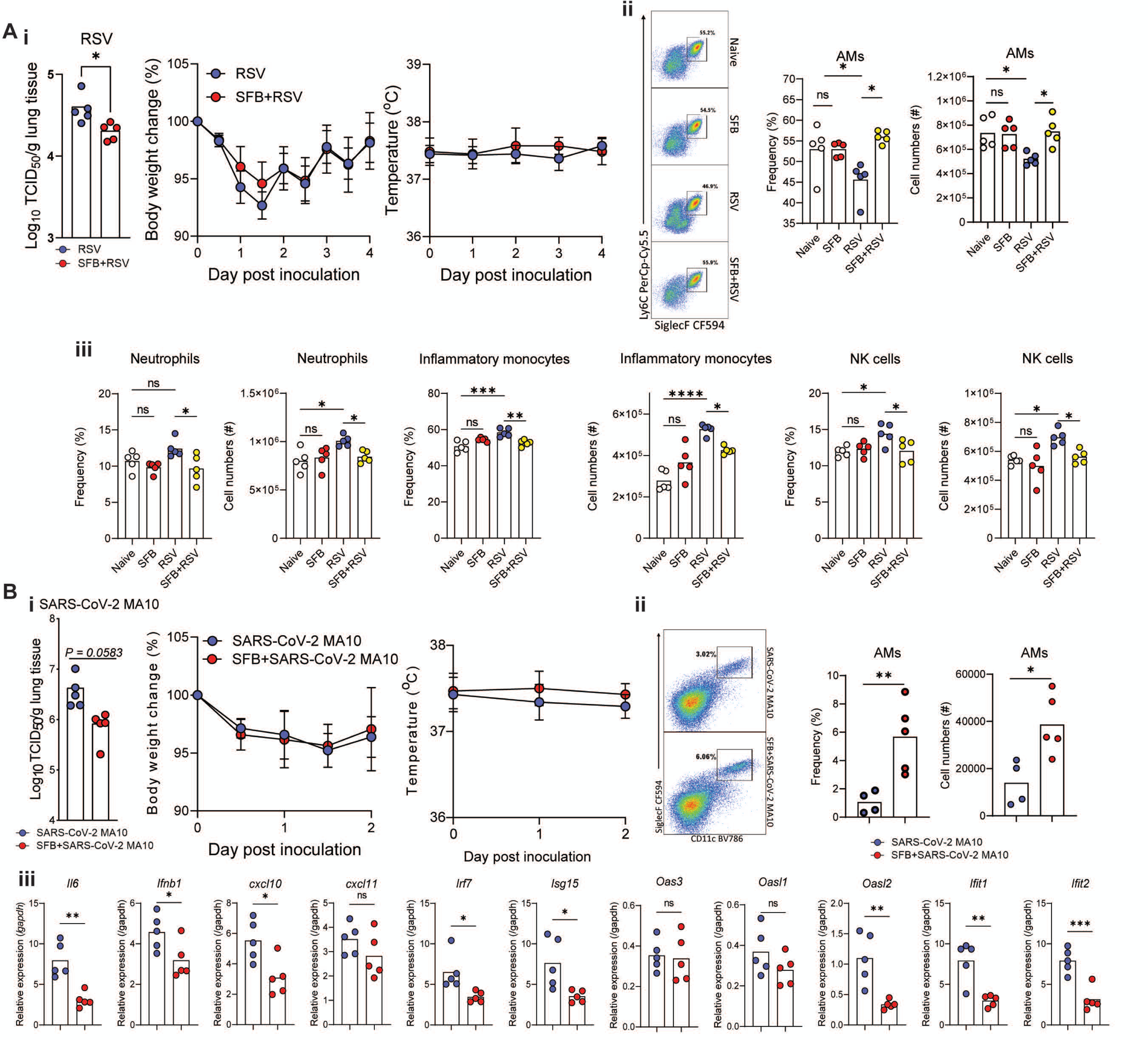

**Figure S5 (related F4).**
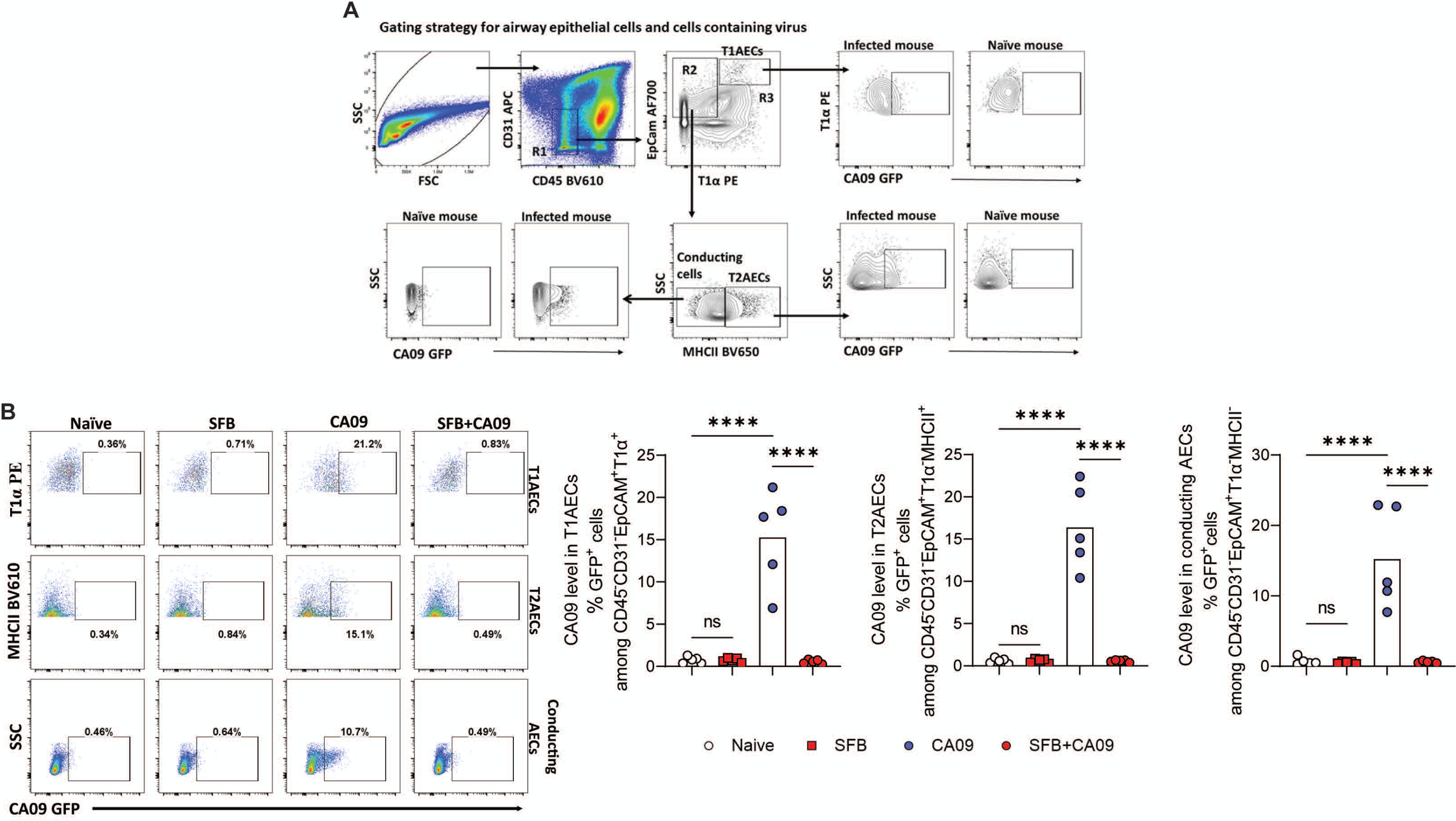

**Figure S6 (related F6).**
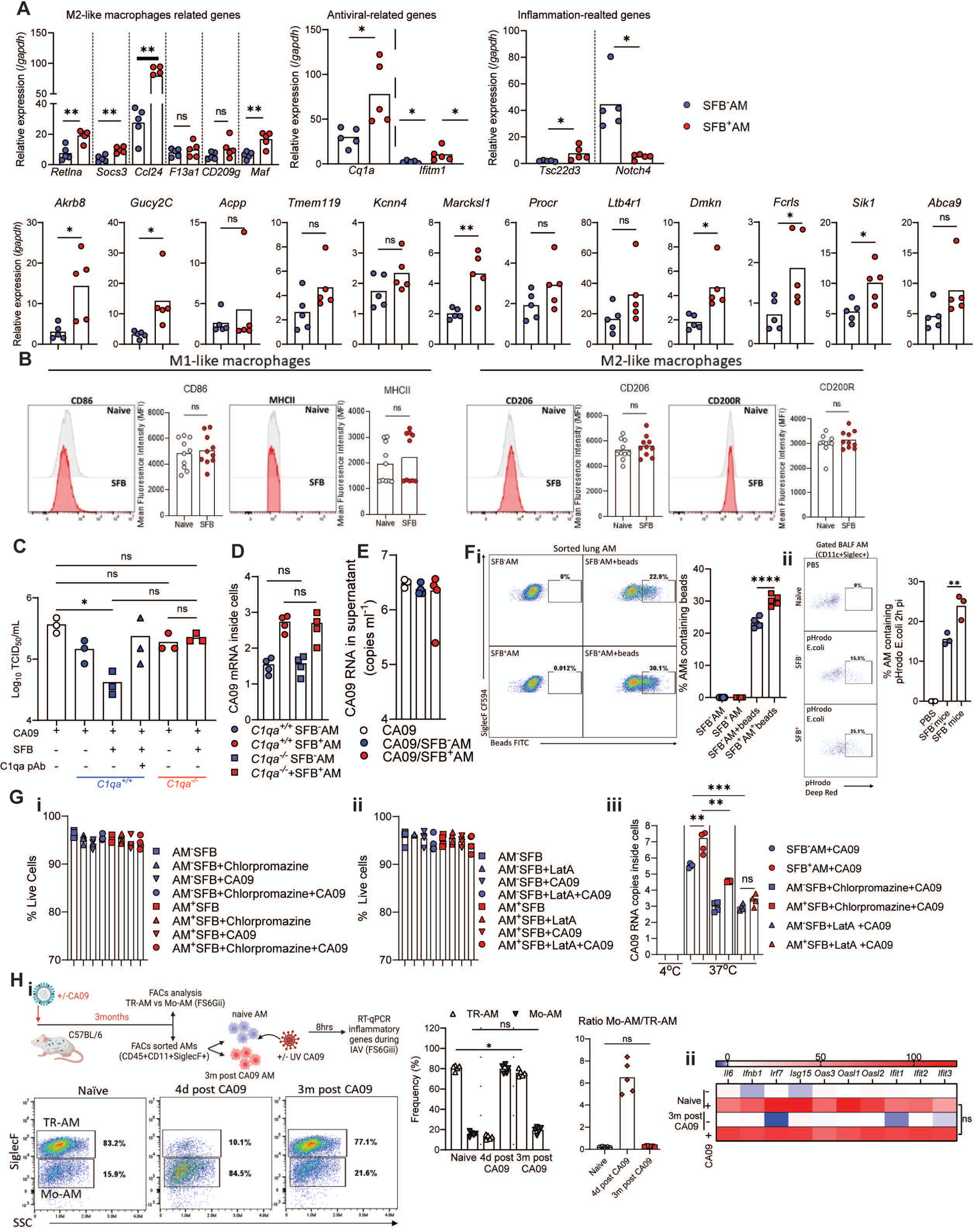

**Figure S7 (related F6 & F7).**
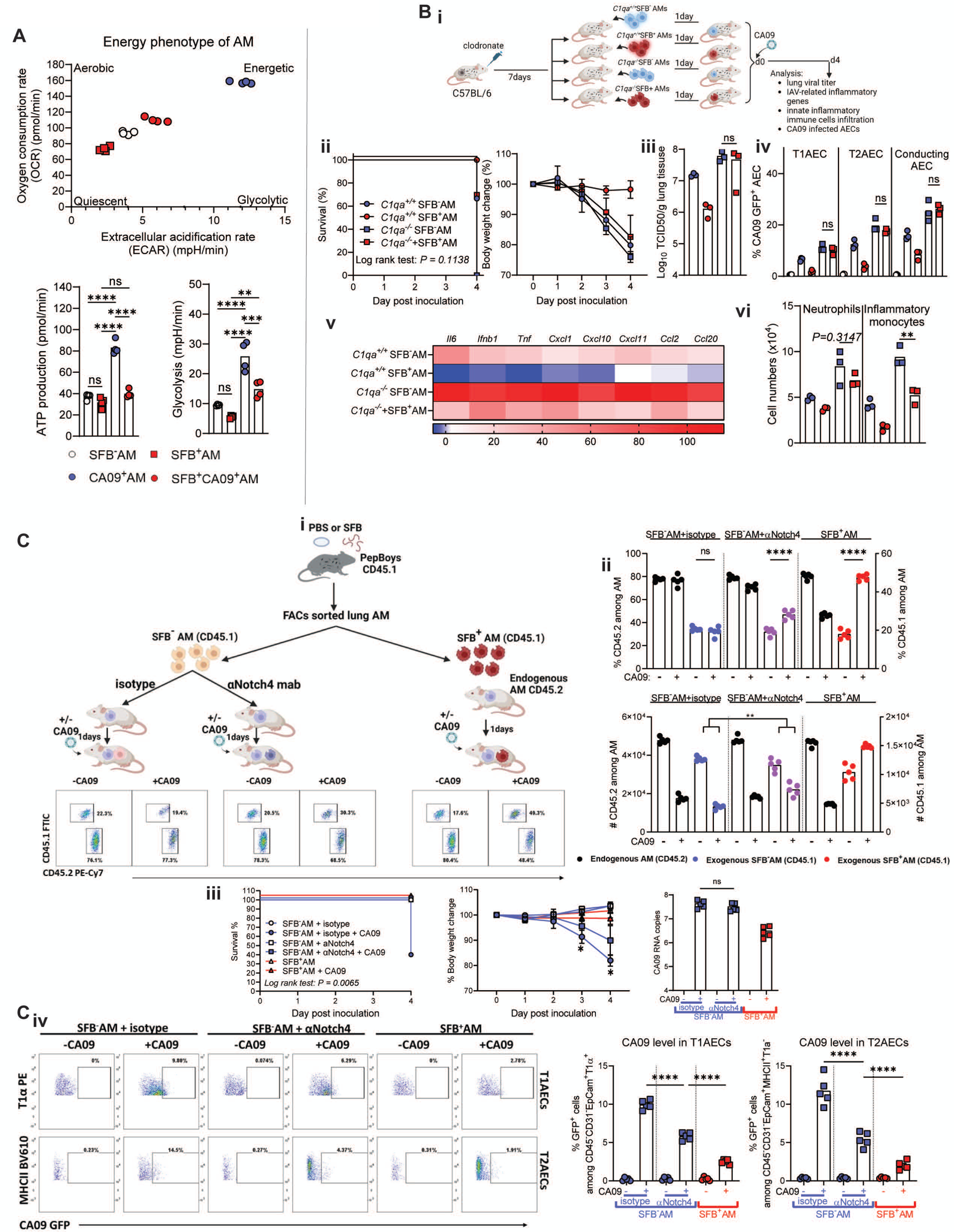

